# SUBCELLULAR FUNCTIONS OF *UBE3A* ISOFORMS DRIVE SYNAPTIC DYSFUNCTION IN ANGELMAN SYNDROME

**DOI:** 10.64898/2026.03.24.713622

**Authors:** Martina Biagioni, Federica Baronchelli, Matteo Monachello, Chiara Ongaro, Edoardo Favriga, Marco Erreni, Alessandra Folci, Davide Pozzi, Matteo Fossati

## Abstract

Genetic defects of the gene encoding the ubiquitin ligase UBE3A cause a severe neurodevelopmental disorder, the Angelman syndrome (AS). The pathophysiology of AS remains unclear, hindering the development of effective therapies. Using AS animal models, we show here that UBE3A controls the development of excitatory and distinct subtypes of inhibitory synapses in cortical pyramidal neurons through cell-autonomous mechanisms, ultimately leading to alteration of synaptic transmission and hyperexcitability. Replacing endogenous *Ube3a* with individual isoforms, we demonstrate that their uneven nuclear (hUBE3A isoform 1) and cytosolic (hUBE3A isoforms 2/3) distribution is critical for regulating distinct aspects of synaptic development. We also define the molecular requirements underpinning this regulation, showing that: (*i*) both nuclear and cytosolic UBE3A rely on their ubiquitin ligase activity to ensure proper assembly of synapses; (*ii*) in addition to the nucleus, UBE3A isoform 1 is also localized in the cytosol, where it is functionally interchangeable with UBE3A isoform 3. Our findings identify a subcellular distribution-dependent mechanism by which UBE3A coordinates cortical circuit development and suggest pathogenic mechanisms of AS.

## INTRODUCTION

Angelman syndrome (AS) is a debilitating neurodevelopmental disorder of genetic origins that affects 1:20,000 individuals. Primary symptoms of AS are severe intellectual disability (ID), developmental delay, ataxic movements, absence of speech, epilepsy, and behavior-specific traits, including unusually happy demeanor (Buiting *et al*, 2016; Williams, 2005). Although different mechanisms underlie AS, the common denominator among all genetic alterations is the loss of the maternally transmitted *UBE3A* gene, which is therefore the causal factor of AS (Kishino *et al*, 1997; Matsuura *et al*, 1997; Judson *et al*, 2014).

*UBE3A* encodes an E3 ubiquitin ligase of the Homologous E6-AP Carboxy Terminus (HECT) family, which catalyzes the conjugation of ubiquitin moieties to its substrates, ultimately targeting them for proteasome-mediated degradation (Mabb & Ehlers, 2010; George *et al*, 2018; Mabb *et al*, 2011). In line with this, defective protein degradation is considered a major determinant of AS pathogenesis (Sell & Margolis, 2015). Importantly, excessive dosage or function of *UBE3A* is strongly associated to autism (Cook *et al*, 1997; Yi *et al*, 2015; Xing *et al*, 2023), further pointing to a crucial role of *UBE3A* in neurodevelopment. How *UBE3A* alterations lead to neuronal dysfunction in these neurodevelopmental disorders remains largely unknown.

*UBE3A* codes for three enzyme isoforms that are generated by alternative splicing and differ in their amino termini (Yamamoto *et al*, 1997). In mature neurons, longer isoforms 2 and 3 (UBE3A-Iso2 and Iso3) are mainly localized in the cytosol, while the shorter isoform 1 (UBE3A-Iso1) lacking an N-terminal extension of 21 amino acids (aa) is enriched in the nucleus and represents ∼80% of total UBE3A (Avagliano Trezza *et al*, 2019; Bossuyt *et al*, 2021; van Esbroeck *et al*, 2025). This molecular diversity and isoform-specific subcellular localization is also conserved in mice (Avagliano Trezza *et al*, 2019; Miao *et al*, 2013; Valluy *et al*, 2015). Interestingly, some missense pathogenic mutations of *UBE3A* are associated with a loss of its nuclear localization (Bossuyt *et al*, 2021). In addition, a recent study generating isoform-specific knock-out mice further supported the notion that UBE3A function in the nucleus is an important determinant for AS pathophysiology (Avagliano Trezza *et al*, 2019). However, patients carrying mutations abrogating UBE3A nuclear localization display milder phenotypes (Sadhwani *et al*, 2018), which indicates that UBE3A cytosolic function cannot be entirely neglected. Hence, it is important to better elucidate UBE3A isoform-specific roles in neuronal function and AS pathophysiology, ultimately identifying AS pathogenic mechanisms and novel therapeutic approaches.

Studies employing cellular and animal models of AS indicate synaptic impairment as a major cellular hallmark. The emerging picture delineates a complex situation, where UBE3A loss has different effects in distinct cell types and neuronal circuits. Decreased density of dendritic spines, where most glutamatergic synapses of the central nervous system reside (Yuste & Bonhoeffer, 2004), is reported in several brain regions, including the cortex, hippocampus and cerebellum (reviewed in (Biagioni *et al*, 2024)). Electrophysiological changes of glutamatergic transmission are widespread in multiple excitatory neurons, such as those in the visual(Wallace *et al*, 2017) and prefrontal cortex (PFC) (Rotaru *et al*, 2018; Sidorov *et al*, 2018), hippocampus (Kaphzan *et al*, 2011, 2013) and dorsomedial striatum (Rotaru *et al*, 2023). UBE3A is also critical to inhibitory connectivity, where dysfunction of GABAergic transmission may underlie the seizure susceptibility observed in AS (Rotaru *et al*, 2018; Wallace *et al*, 2012; Judson *et al*, 2016). Notwithstanding, whether and how *UBE3A* defects impact the development and specification of distinct inhibitory synapse subtypes remains elusive. Additionally, the contribution of UBE3A isoforms to the assembly of synaptic connectivity is not known, and its understanding might uncover novel pathogenic mechanisms underlying cognitive and behavioral alterations in AS.

In the present study, we use sparse in utero electroporation (IUE) of layer 2/3 pyramidal neurons of the mouse somatosensory cortex to dissect the role of UBE3A isoforms at the single-cell level in vivo. We first demonstrate that the inactivation of *Ube3a* during cortical development impairs the formation of excitatory and perisomatic inhibitory synapses, and the maturation of GABAergic synapses of the axon initial segment (AIS) through cell-autonomous mechanisms. By replacing endogenous *Ube3a* with individual human isoforms in utero, we show that the subcellular localization of UBE3A critically regulates distinct aspects of synaptic development: whereas nuclear UBE3A rescues dendritic spine formation and maturation of AIS inhibitory synapses, the cytosolic enzyme regulates perisomatic inhibitory connections. Importantly, similar synaptic impairments are also detected in the most used mouse model of AS (Jiang *et al*, 1998) and rescued by re-expression of single isoforms, thus enhancing the translational relevance of these findings. Together, our data uncover the crucial role of UBE3A, and its subcellular localization, in the specification and maturation of distinct subtypes of excitatory and inhibitory synapses, with implications for the pathogenesis of AS.

## RESULTS

### Loss of *Ube3a* impairs the formation of excitatory synapses

To investigate the role of UBE3A in synaptic development in vivo, we employed IUE - a method of choice to study synaptogenesis at single-cell resolution (Fossati *et al*, 2019, 2016, 2022) - at embryonic day (E)15.5 to target layer 2/3 cortical pyramidal neurons (CPNs) of the mouse somato-sensory cortex in their native environment (Fig 1A). Endogenous *Ube3a* was inactivated using the Clustered Regularly Interspaced Short Palindromic Repeats (CRISPR)/Cas9 system, which showed more than 90% efficiency in vivo as indicated by the absence of UBE3A signal in CPNs expressing the gRNA against *Ube3a* (Figure S1). The co-expression of the myristoylated fluorescent reporter mVenus was used to visualize neuronal morphology and analyze dendritic spines as proxy of glutamatergic synapses (Fig 1B) (Yuste & Bonhoeffer, 2004). We found that UBE3A depletion significantly reduced the density of dendritic spines in juvenile mice (post-natal day, P21) (Fig 1C; Figure S2), without affecting their morphology using either confocal or super-resolution stimulated emission depletion (STED) microscopy (Fig 1D-F; Figure S3). To directly monitor excitatory synapses, EGFP-tagged fibronectin intrabodies generated with mRNA display (FingR) against PSD-95 (Gross *et al*, 2013; Fossati *et al*, 2022), the major scaffolding protein of excitatory synapses (Sheng & Kim, 2011), were expressed in utero (Fig 1G). Consistent with the observed alterations of spine number, the density of PSD-95 clusters decreased to 78% ± 7% of the control value, without variations in their size and fluorescence intensity (Fig 1H-J; Figure S2). We also found similar impairments in neurons knockdown (KD) for *Ube3a* using a short hairpin RNA (shRNA) (Figure S2), further supporting the role of UBE3A in excitatory synaptogenesis.

**Fig 1.**
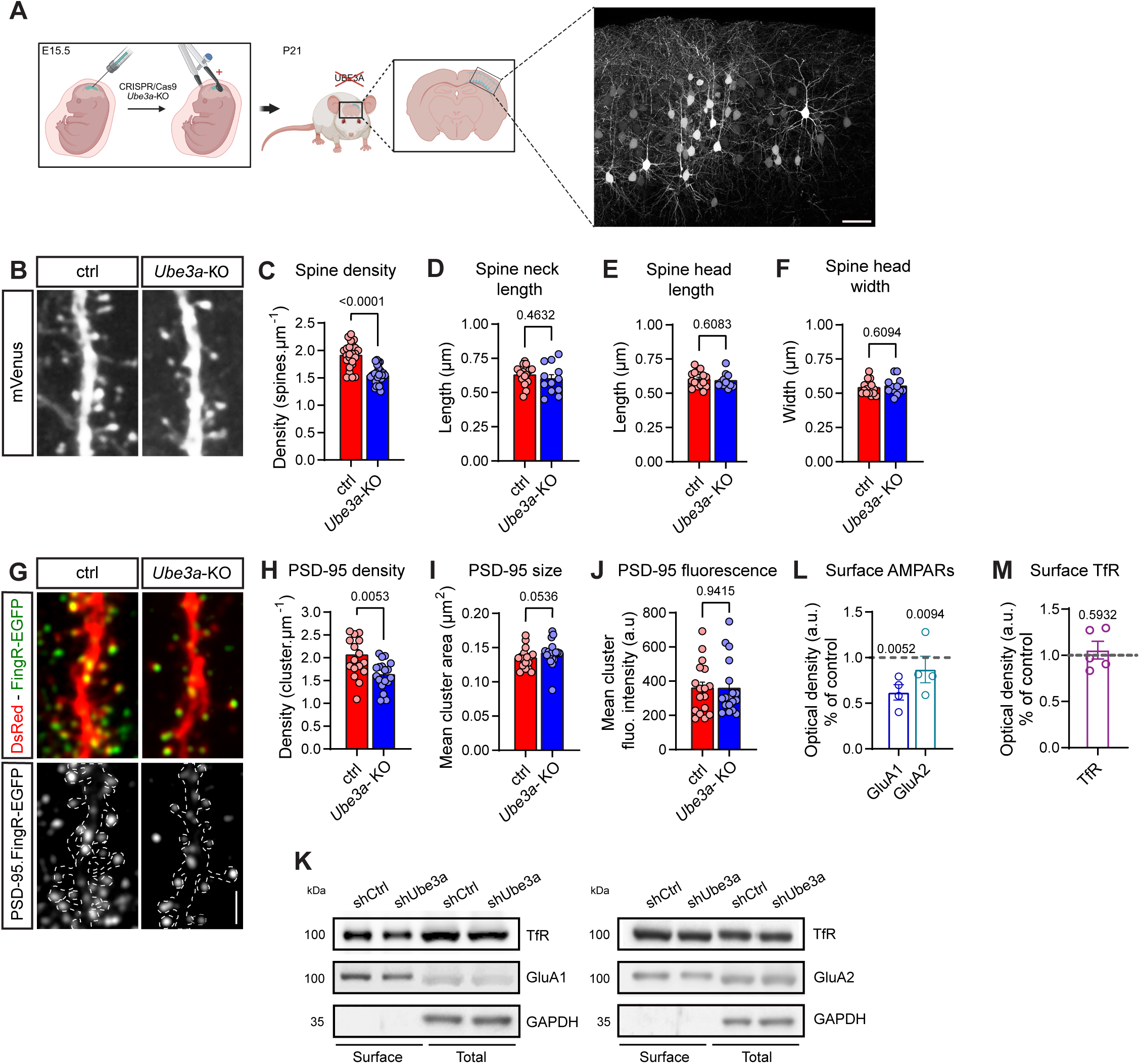
*Ube3a* inactivation impairs excitatory synaptogenesis. **(A)** Schematic (left) and representative image (right) of sparse labeling of layer 2/3 CPNs after in utero electroporation (IUE) with mVenus. E15.5, embryonic day 15.5; P21: postnatal day 21. Scale bar: 50µm. Created in https://BioRender.com. **(B)** Segments of dendrites expressing control CRISPR/Cas9 (ctrl) or against *Ube3a* (*Ube3a*-KO) along with mVenus to visualize dendritic spines in juvenile mice. Scale bar: 2 µm. **(C-F)** Quantification of density (C), neck length (D), head length (E) and width (F) of dendritic spines in juvenile mice. Spine density analysis (C): n_ctrl_ = 23 (6); n*_Ube3a_*_-KO_ = 28 (7); other parameters analysis (D-F): n_ctrl_ = 15 (6); n*_Ube3a_*_-KO_ = 12 (3). **(G)** Segments of dendrites illustrating the effect of Ube3a inactivation on PSD-95 clusters. Dashed lines (bottom) mark the contours defined by the cytosolic filler DsRed. Scale bar: 2 µm. **(H-J)** Quantification of density (H), size (I) and fluorescence intensity (J) of PSD-95 clusters in juvenile mice. Cluster density analysis: n_ctrl_ = 31 (5); n*_Ube3a_*_-KO_ = 24 (6); cluster area and fluorescence intensity analysis: n_ctrl_ = 17 (3); n*_Ube3a_*_-KO_ = 18 (3). **(K)** Cell surface expression of GluA1 (left) and GluA2 (right) subunits of AMPA receptors (AMPARs) in DIV17 primary cultures of cortical neurons from E18.5 mouse embryos infected with lentiviral vectors driving the expression of shControl (shCtrl) or sh*Ube3a*. GAPDH and Transferrin Receptor (TfR) are used as loading control and to check efficient and homogenous isolation of surface proteins, respectively. **(L-M)** Normalized cell surface abundance of GluA1 and GluA2 (L) and TfR (M) as shown in (K). Dotted line indicates control values. n = 4. Statistics: bars indicate mean ± SEM. Numbers in parentheses indicate the number of animals. p values are indicated in the graphs. Student t-test or Mann-Whitney test.

Given the crucial role of PSD-95 as binding platform to regulate the abundance of glutamatergic receptors at synapses, a major determinant of synaptic strength (Opazo *et al*, 2012), we assessed the number of α-amino-3-hydroxy-5-methyl-4-isoxazolepropionic acid receptors (AMPARs) expressed at the cell surface in primary cultures of cortical neurons transduced with Lentiviral particles driving the expression of shRNAs against *Ube3a* (Figure S1). Using cell surface biotinylation, we found that the abundance of GluA1-and GluA2-containing AMPARs, but not that of transferrin receptors, is significantly decreased in *Ube3a*-KD neurons (Fig 1K-M). Together, these results indicate that the loss of *Ube3a* during neurodevelopment alters the assembly of excitatory synapses in the neocortex of juvenile mice.

### Loss of *Ube3a* affects the development of distinct subclasses of inhibitory synapses onto CPNs

Distinct classes of interneurons possess specific molecular programs underlying their unique and stereotyped connectivity onto CPNs (Tremblay *et al*, 2016; Favuzzi *et al*, 2019). A handful of studies suggested that inhibitory synaptic connections are affected in AS animal models (Wallace *et al*, 2012; Gu *et al*, 2018; Judson *et al*, 2016; Rotaru *et al*, 2018). However, whether UBE3A operates in distinct subtypes of inhibitory synapses remains elusive. To address this issue, we in utero electroporated fluorescent FingRs recognizing gephyrin (GPHN) (Gross *et al*, 2013; Fossati *et al*, 2019, 2022), the major scaffolding protein of inhibitory synapses (Tyagarajan & Fritschy, 2014) together with CRISPR/Cas9 to ablate *Ube3a* (Fig 2A). GPHN clusters were analyzed by high-resolution confocal microscopy in distal dendrites, perisomatic area and AIS, the three subcellular compartments where major subclasses of cortical interneurons synapse onto CPNs, namely somatostatin-positive cells (STT+) and parvalbumin-positive (PV+) basket and chandelier cells, respectively (Tremblay *et al*, 2016). In juvenile mice, UBE3A depletion using CRISPR/Cas9 altered neither the density (Fig 2B) nor other morphometric parameters reflecting the maturation and complexity of dendritic inhibitory synapses, such as the size, fluorescent intensity and localization of GPHN clusters (Fig 2C-E) (Fossati *et al*, 2016; Chen *et al*, 2012). We then analyzed perisomatic synapses. FingR fluorescence reliably matches the expression levels of the endogenous targets due to its transcriptional regulation system, which however causes the accumulation of FingR molecules in the nucleus (Gross *et al*, 2013). To ensure that FingR-positive puncta visualized in the perisomatic region corresponded to inhibitory synapses, we assessed only the number of GPHN clusters co-localizing with vGAT-positive puncta through somatic volume (Fig 2F-G). We found significant alterations in the formation of perisomatic inhibitory synapses in *Ube3a*-KO neurons, which showed a reduction to 81% ± 0.3% compared to control neurons (Fig 2F-H). Similar to PSD-95, GPHN contains binding sites for subunits of type A γ -aminobutyric acid receptors (GABA_A_Rs), ultimately regulating their number in the postsynaptic membrane (Tretter *et al*, 2012). Consistent with a reduced density of perisomatic GPHN clusters, we assessed the surface expression of β3-containing GABA_A_Rs in dissociated cortical neurons and found that it is significantly decreased in *Ube3a*-KD neurons compared to cells expressing shControl (Fig 2I-J).

**Fig 2.**
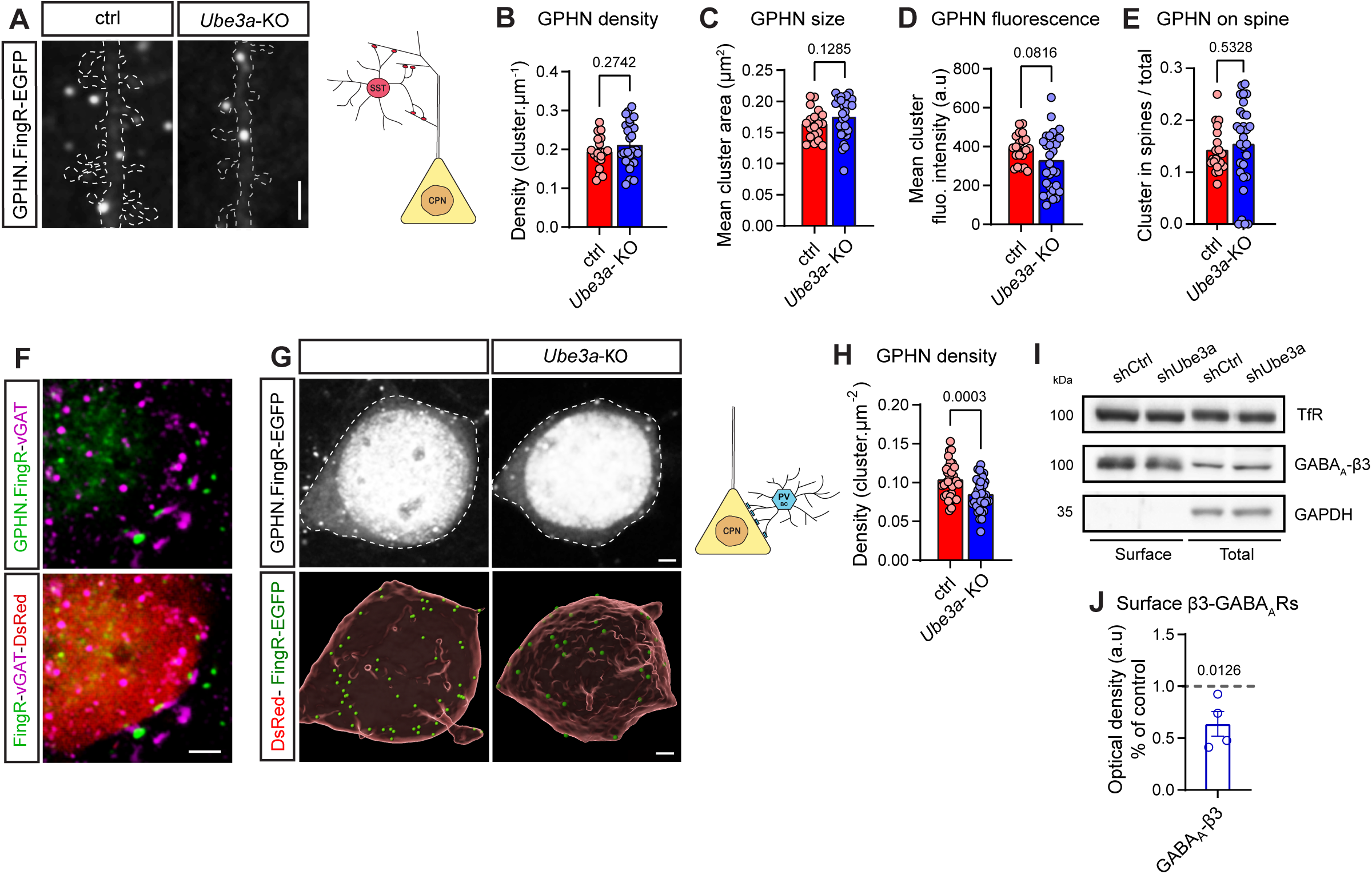
Selective regulation of perisomatic inhibitory connectivity by UBE3A. **(A)** Left: segments of dendrites expressing control CRISPR/Cas9 (ctrl) or against *Ube3a* (*Ube3a*-KO) along with GPHN.FingR-EGFP in juvenile mice. Dashed lines indicate the contours defined by the cytosolic filler DsRed. Scale bar: 2 µm. Right: schematic showing inhibitory innervation onto distal dendrites of CPNs made by Somatostatin-positive cortical interneurons (SST). **(B-E)** Quantification of density (B), size (C), fluorescence intensity (D) and compartmentalization (E) of GPHN clusters on distal dendrites of electroporated CPNs in juvenile mice. n_ctrl_ = 20 (3); n*_Ube3a-_*_KO_ = 27 (4). **(F)** Representative image showing the apposition between GPHN clusters and vGAT-positive presynaptic puncta on the somatic surface of electroporated CPNs. Scale bar: 2 µm. **(G)** Left: representative images of inhibitory synapses identified using GPHN.FingR-EGFP onto the soma of ctrl or *Ube3a*-KO neurons (top), and corresponding 3D reconstructions showing GPHN clusters (green) on top of somatic surfaces (red). Dashed lines (top) delineate the contours of neuronal somata defined by the cytosolic filler DsRed. Scale bars: 2 µm. Right: schematic showing inhibitory innervation onto the soma of CPNs made by Parvalbumin-positive basket cells (PVBCs). **(H)** Quantification of perisomatic GPHN cluster density in juvenile mice. n_ctrl_ = 36 (6); n*_Ube3a-_*_KO_ = 36 (6). **(I)** Cell surface expression of β3-containing GABAARs in DIV17 cortical neurons obtained from E18.5 mouse embryos and transduced with lentiviral vectors driving the expression of shControl (shCtrl) or sh*Ube3a*. GAPDH and Transferrin Receptor (TfR) are used as loading control and to test efficient and homogenous enrichment of surface proteins. Quantification of surface TfR is already shown in Fig. 1M. **(J)** Normalized cell surface abundance of GABA_A_R- β3 as shown in (I). Dotted line indicates control values. n = 4. Statistics: bars indicate mean ± SEM. Numbers in parentheses indicate the number of animals. p values are indicated in the graphs. Student t-test with Welch’s correction.

To identify AIS inhibitory synapses, brain slices obtained from electroporated animals at P21 were immunostained for ankyrin G (AnkG) protein, which is selectively expressed in the AIS (Ogawa & Rasband, 2008) (Fig 3A). The density and size of these synapses were not affected by the loss of *Ube3a* (Fig 3B-C). However, we observed a significant reduction in the fluorescence intensity of AIS GPHN clusters, indicating that GPNH recruitment at nascent synapses was impaired in *Ube3a*-KO neurons (Fig 3D). To test whether this defective accumulation of GPHN resulted in changes of the abundance of synaptic GABA_A_Rs, we used neuronal cultures and carried out live-cell staining of surface GABA_A_Rs using antibodies recognizing extracellular epitopes of different receptor subunits. First, we confirmed that a defective recruitment of AIS-localized GPHN clusters was recapitulated in dissociated cortical neurons infected with lentiviral particles to knock-down *Ube3a* (Fig 3E). In agreement with the role of the GPHN hexagonal lattice in stabilizing GABA_A_Rs (Tyagarajan & Fritschy, 2014), the abundance of both α2- and γ 2-containing GABA_A_Rs was reduced in *Ube3a*-KD neurons (Fig 3F-I). Notably, the change was more pronounced for the α2 subunit (Fig 3F-G), which contains a GPHN-binding motif and which is specifically enriched at AIS inhibitory synapses compared to γ2 (Fig 3H-I); the latter is instead a ubiquitous component of most GABA_A_ pentamers and does not directly interact with GPHN (Luscher *et al*, 2011; Panzanelli *et al*, 2011).

**Fig 3.**
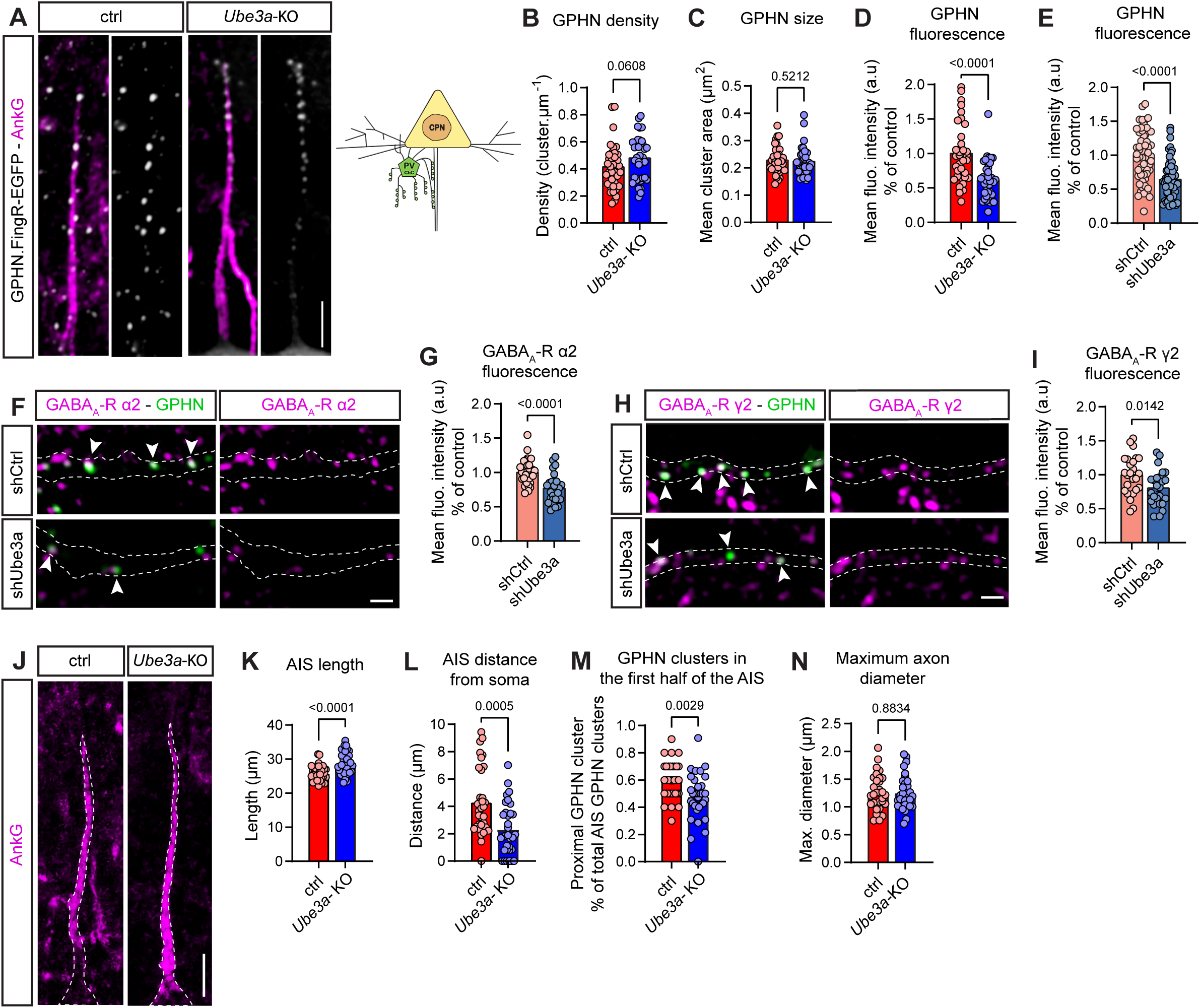
*Ube3a* loss alters AIS structural organization and axo-axonic inhibitory innervation. **(A)** Left: segments of dendrites of ctrl and *Ube3a*-KO neurons expressing GPHN.FingR-EGFP to label inhibitory synapses and stained for AnkG to identify AIS. Scale bar: 5 µm. Right: schematic showing AIS inhibitory innervation of CPNs made by Parvalbumin-positive chandelier cells (PVChCs). **(B-D)** Quantification of density (B), size (C) and normalized fluorescence intensity (D) of AIS GPHN clusters in juvenile mice. n_ctrl_ = 36 (4); n*_Ube3a_*_-KO_ = 36 (5). **(E)** Quantification of normalized fluorescence intensity of AIS-localized GPHN clusters in control or *Ube3a* knocked-down cortical neurons at DIV17. n_shCtrl_ = 58 (6); n_shUbe3a_ = 56 (6). **(F, H)** Live-cell staining of surface α2- (F) and γ2-containing GABA_A_Rs (H) at AIS in DIV17 cortical neurons co-expressing shControl or sh*Ube3a* and GPHN.FingR-EGFP. Arrowheads indicate association between GABA_A_Rs and GPHN clusters. Dashed lines delineate the contours of AIS defined by AnkG staining (not shown). Scale bar: 2 µm. **(G, I)** Quantification of normalized fluorescence intensity of AIS-localized α2- (G) and γ2-containing GABAARs (I) clusters in control or *Ube3a* knocked-down cortical neurons at DIV17. α2-GABA_A_R: n_shCtrl_ = 32 (4); n_shUbe3a_ = 29 (4); γ2-GABA_A_R: n_shCtrl_ = 25 (3); n_shUbe3a_ = 25 (3). **(J)** Representative images illustrating the effect of *Ube3a* inactivation on AIS length and point of origin. Dashed lines indicate the contours defined by the cytosolic filler DsRed. Scale bar: 5 µm. **(K-N)** Quantification of AIS length (K), distance between AIS origin and neuronal soma (L), percentage of GPHN clusters located in the first half of the AIS (M) and maximum axon diameter (N). n_ctrl_ = 34 (4); n_Ube3a-KO_ = 33 (5). Statistics: bars indicate mean ± SEM. Numbers in parentheses indicate the number of animals or neuronal cultures. p values are indicated in the graphs. Student t-test with Welch’s correction or Mann-Whitney test.

The AIS is a critical site for the integration of synaptic inputs and action potential initiation and undergoes homeostatic changes to finely tune neuronal excitability (Wefelmeyer *et al*, 2015; Pan-Vazquez *et al*, 2020; Grubb & Burrone, 2010; Evans *et al*, 2013). Alterations of its molecular composition and function were suggested to contribute to neuronal hyperexcitability in AS (Rayi & Kaphzan, 2021; Kaphzan *et al*, 2013, 2011). Therefore, we analyzed AIS morphometric features, including its length, diameter and position (Wefelmeyer *et al*, 2015). We found that *Ube3a* deletion did not affect axonal diameter of the AIS (Fig 3N) but significantly increased its length and its point of origin, which was shifted closer to neuronal soma (Fig 3J-L). AIS relocation also caused a mismatch with the position of inhibitory synapses (Wefelmeyer *et al*, 2015), which were less numerous in the first part of the AIS (Fig 3M). Taken together, these data revealed defects in distinct aspects of the development of specific subtypes of inhibitory synapses onto CPNs, indicating that the loss of *Ube3a* impairs the assembly of inhibitory connectivity in the mouse cortex.

### UBE3A controls excitatory and inhibitory synaptic transmission

Given the results obtained with our morphometric analysis of excitatory and inhibitory synapses, we tested the physiological consequences of *Ube3a* inactivation on synaptic transmission. To this end, we carried out whole-cell patch clamp recording in acute brain slices obtained from juvenile mice electroporated in utero with either control CRISPR/Cas9 or against endogenous *Ube3a* (Figure S1). The co-electroporation of GFP as cytosolic filler allowed the identification of transfected neurons (Fig 4A). Miniature excitatory and inhibitory postsynaptic currents (mEPSCs and mIPSCs, respectively) were simultaneously recorded on the same neuron to provide a measurement of the excitatory/inhibitory (E/I) balance. In *Ube3a*-KO brains, the frequency of mEPSCs was significantly decreased, without any significant change in their amplitude (Fig 4B-D). This observation is in line with a decreased number of excitatory synapses (Fig 1C, J; Figure S2). In addition, the frequency of mIPSCs was also significantly lower in *Ube3a*-KO neurons. This result is in accordance with an impairment of inhibitory synapses that are located close to the soma of CPNs, where inhibitory currents were recorded (Tremblay *et al*, 2016) (Fig 4E-G). Strikingly, when we calculated the ratio between synaptic excitation and inhibition, we found a significant shift toward increased excitation (Fig 4H). Together, these results indicate that the loss of UBE3A alters synaptic development, which is also accompanied by changes in functional organization of the AIS, ultimately impairing synaptic transmission and the excitation-to-inhibition ratio in AS.

**Fig 4.**
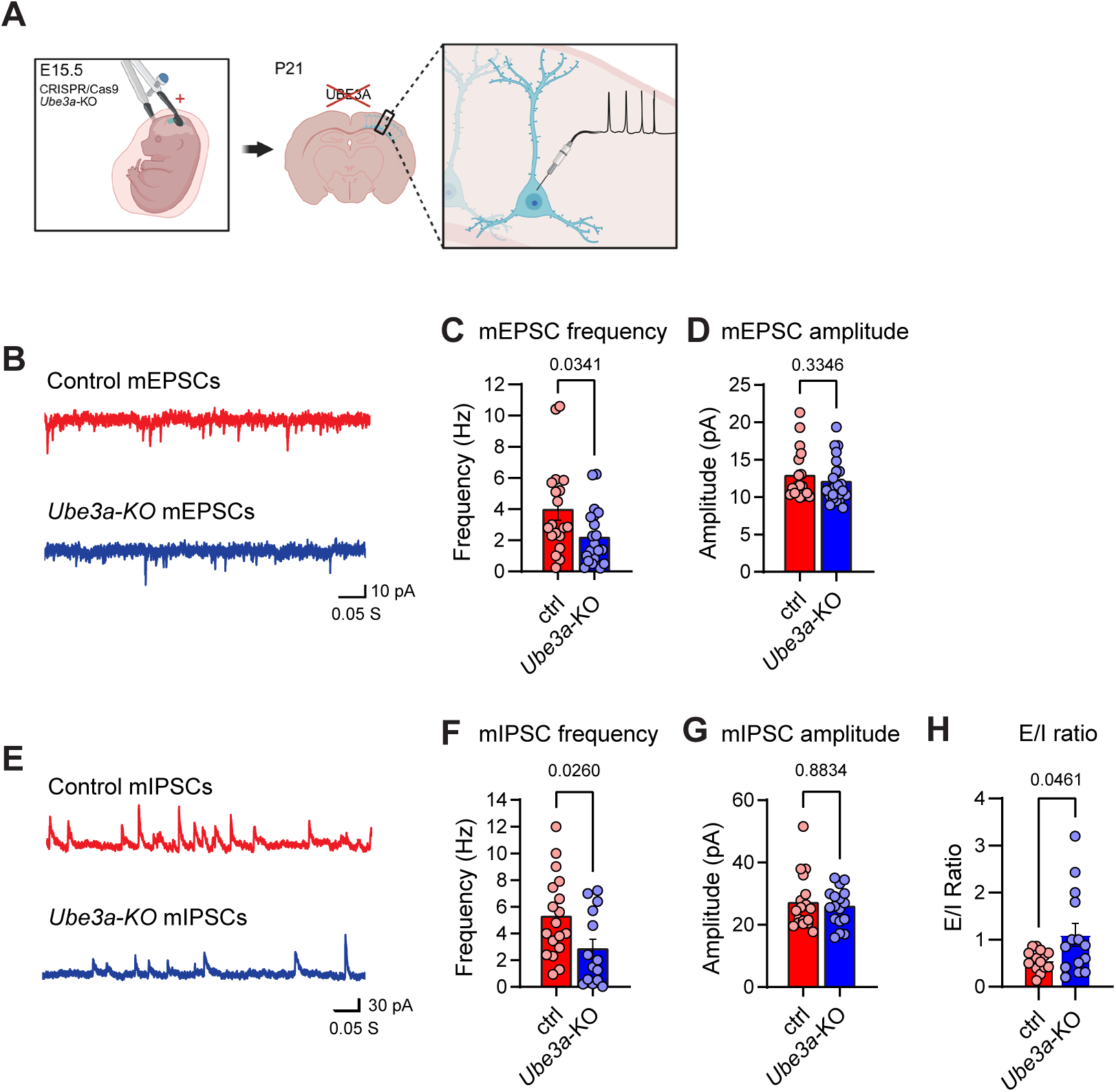
*Ube3a* loss alters excitatory and inhibitory synaptic transmission and impairs E/I ratio towards hyperexcitability. **(A)** Schematic illustrating patch-clamp recording of electroporated layer 2/3 CPNs co-expressing control CRISPR/Cas9 (ctrl) or against *Ube3a* (*Ube3a*-KO) and GFP from acute brain slices of juvenile mice. Created in https://BioRender.com. **(B, E)** Representative traces of mEPSCs (B) and mIPSCs (E) in control and *Ube3a*-KO electroporated neurons. **(C-D, F-G)** Quantification of mEPSC and mIPSC frequency (C, F) and amplitude (D, G) in control and *Ube3a*-KO electroporated neurons. mEPSCs: n_ctrl_ = 20 (4); n*_Ube3a_*_-KO_ = 19 (4); mIPSC: n_ctrl_ = 18 (4); n_Ube3a-KO_ = 14 (4). **(H)** Quantification of excitatory-inhibitory (E/I) ratio in control and *Ube3a*-KO electroporated neurons. n_ctrl_ = 18 (4); n*_Ube3a_*_-KO_ = 14 (4). Statistics: bars indicate mean ± SEM. Numbers in parentheses indicate the number of animals. p values are indicated in the graphs. Student t-test and Mann-Whitney test.

### Nuclear and cytosolic isoforms of UBE3A are crucial for synapse development

Recent work suggested that different UBE3A isoforms generated by alternative splicing are distributed in distinct intracellular compartments, the nucleus and cytosol. Yet, their functional relevance to neuronal development and AS pathophysiology is poorly characterized (Avagliano Trezza *et al*, 2019; Sirois *et al*, 2020; Judson *et al*, 2021; Miao *et al*, 2013). To directly test whether the molecular diversity of UBE3A plays a role in synapse assembly and specification, we employed IUE to replace endogenous *Ube3a* with individual human isoforms, which are not targeted by the gRNA against mouse *Ube3a*. Of the three splicing variants of *UBE3A*, we focused on *UBE3A-Iso1* and *UBE3A-Iso3* because: (1) they are the most abundantly expressed, constituting ∼98% of total UBE3A (Judson *et al*, 2021); (2) they are homologous to protein-encoding *mUbe3a-Iso3* and *Iso2*, respectively (Avagliano Trezza *et al*, 2019) (Fig 5A). Using immunostaining and confocal imaging, we first checked that the shorter UBE3A-Iso1 was enriched in the nucleus, while the longer UBE3A-Iso3 was mainly cytosolic in both dissociated cortical neurons and in layer 2/3 CPNs (Fig 5B; Fig 6E) (Avagliano Trezza *et al*, 2019; van Esbroeck *et al*, 2025). In juvenile mice, the selective reinstatement of *UBE3A-Iso1*, but not of *UBE3A-Iso3*, was sufficient to restore the proper number of dendritic spines, suggesting that UBE3A in the nucleus is critical to regulate the formation of excitatory synapses (Fig 5C-D). We obtained similar results when we examined inhibitory synapses of the AIS. The mean fluorescent intensity of GPHN clusters was brought back to control values only in neurons expressing *UBE3A-Iso1* (Fig 5E-F). Next, we assessed the role of *UBE3A* splicing variants in the formation of perisomatic inhibitory synapses. Surprisingly, selective expression of both isoforms led to equivalent results, allowing to rescue the correct density of GPHN clusters in the perisomatic region of CPNs from P21 brains (Fig 5G-H). Collectively, these results indicated that the molecular diversity of UBE3A is a critical determinant of synapse development. Notably, UBE3A-Iso1, which is enriched in the nuclear compartment employing immunocytochemistry, regulates the formation of excitatory synaptic inputs onto layer 2/3 CPNs and the maturation of AIS inhibitory synapses. Instead, the formation of inhibitory synapses onto the somatic region of CPNs was restored upon reinstatement of both isoforms individually, suggesting at least partial functional redundancy.

**Fig 5.**
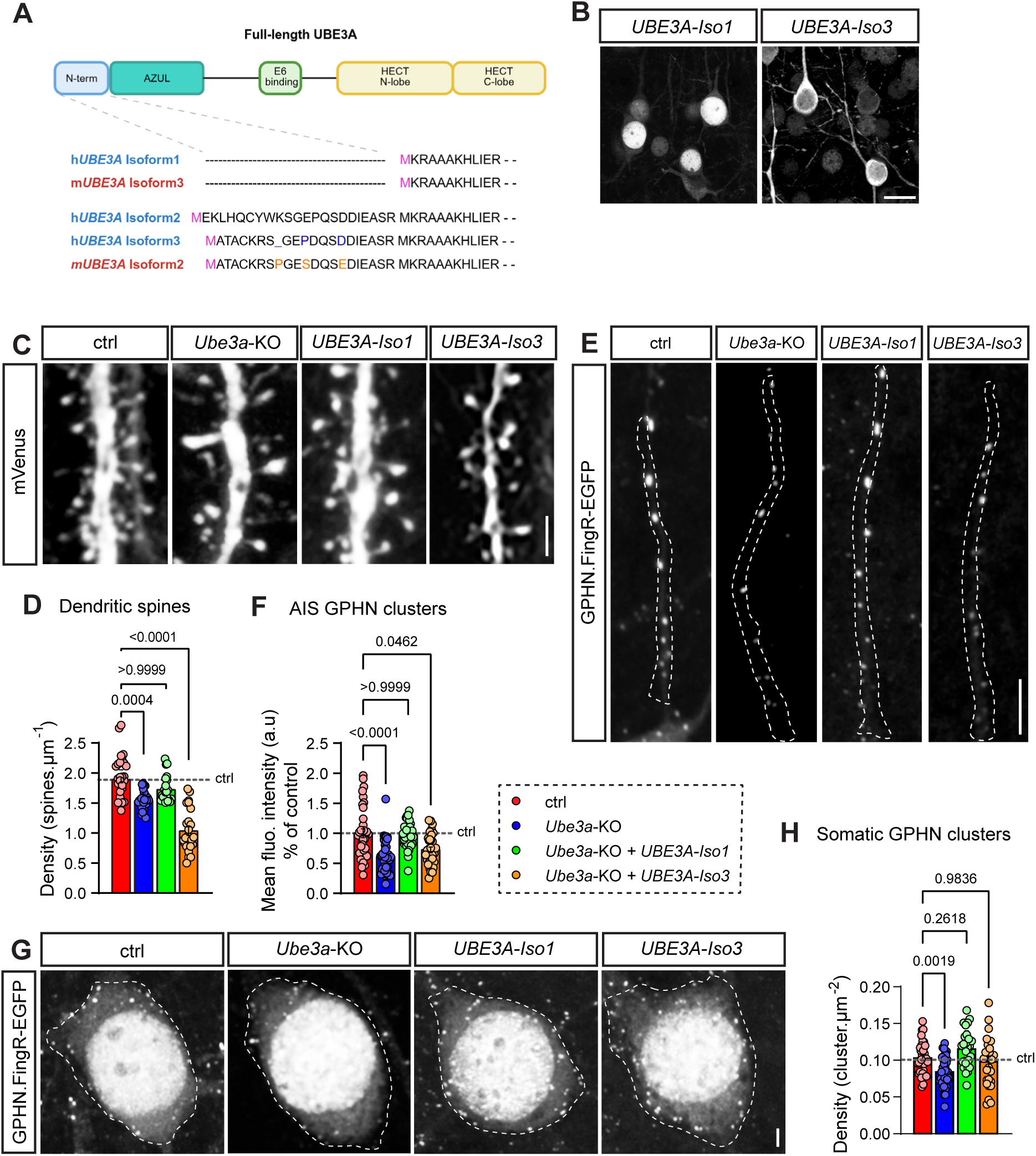
Individual UBE3A isoforms regulate distinct aspects of synaptic development. **(A)** Schematic of UBE3A functional domains (top) and N-terminal amino acidic sequences of human and mouse isoforms (bottom). M residues (pink) represent the initiator Methionine of each isoform and residues in orange highlight differences between murine isoform 2 and the corresponding human isoforms. Dot lines indicate the N-terminal aminoacidic stretch that is absent in shorter human and mouse isoforms 1 and 3, respectively. Created in https://BioRender.com. **(B)** Representative confocal images showing the subcellular distribution of EGFP-tagged UBE3A-Iso1 and Iso3 in electroporated CPNs at P21. Scale bar: 20 µm. **(C)** Dendritic spines in representative segments of oblique dendrites in control (ctrl) or *Ube3a*-KO neurons, or after in utero replacement of endogenous Ube3a with indicated isoforms in juvenile mice. Scale bar: 2 μm. **(D)** Quantification of density of dendritic spines in conditions shown in (C). Dotted line indicates control value. n_ctrl_ = 23 (6); n*_Ube3a_*_-KO_ = 28 (7); n*_UBE3A-Iso1_* = 20 (4); n*_UBE3A-Iso3_* = 23 (5). **(E)** Images of GPHN.FingR-EGFP positive clusters located at AIS in control (ctrl) or *Ube3a*-KO neurons, or after in utero replacement of endogenous Ube3a with indicated isoforms in juvenile mice. Dashed lines define the contours of AnkG fluorescence. Scale bar: 5 μm. **(F)** Quantification of normalized fluorescence intensity of AIS GPHN clusters. Dotted line indicates control value. n_ctrl_ = 36 (4); n*_Ube3a_*_-KO_ = 36 (5); n*_UBE3A-Iso1_* = 38 (3); n*_UBE3A-Iso3_* = 31 (5). **(G)** Perisomatic inhibitory synapses in control (ctrl) and *Ube3a*-KO neurons or after in utero replacement of endogenous *Ube3a* with indicated isoforms in juvenile mice. Dashed lines define the contours of DsRed (ctrl, *Ube3a*-KO) or tdTomato (UBE3A-Iso1, UBE3A-Iso3) fluorescence. Scale bar: 2 μm. **(H)** Quantification of GPHN clusters density in conditions indicated in (G). Dotted line indicates control value. n_ctrl_ = 36 (6); n*_Ube3a_*_-KO_ = 36 (6); n*_UBE3A-_*_Iso1_ = 28 (3); n*_UBE3A_*_-Iso3_ = 26 (3). Statistics: bars indicate mean ± SEM. Numbers in parentheses indicate the number of animals. p values are indicated in the graphs. Welch and Brown-Forsythe one-way ANOVA or Kruskal-Wallis test.

**Fig 6.**
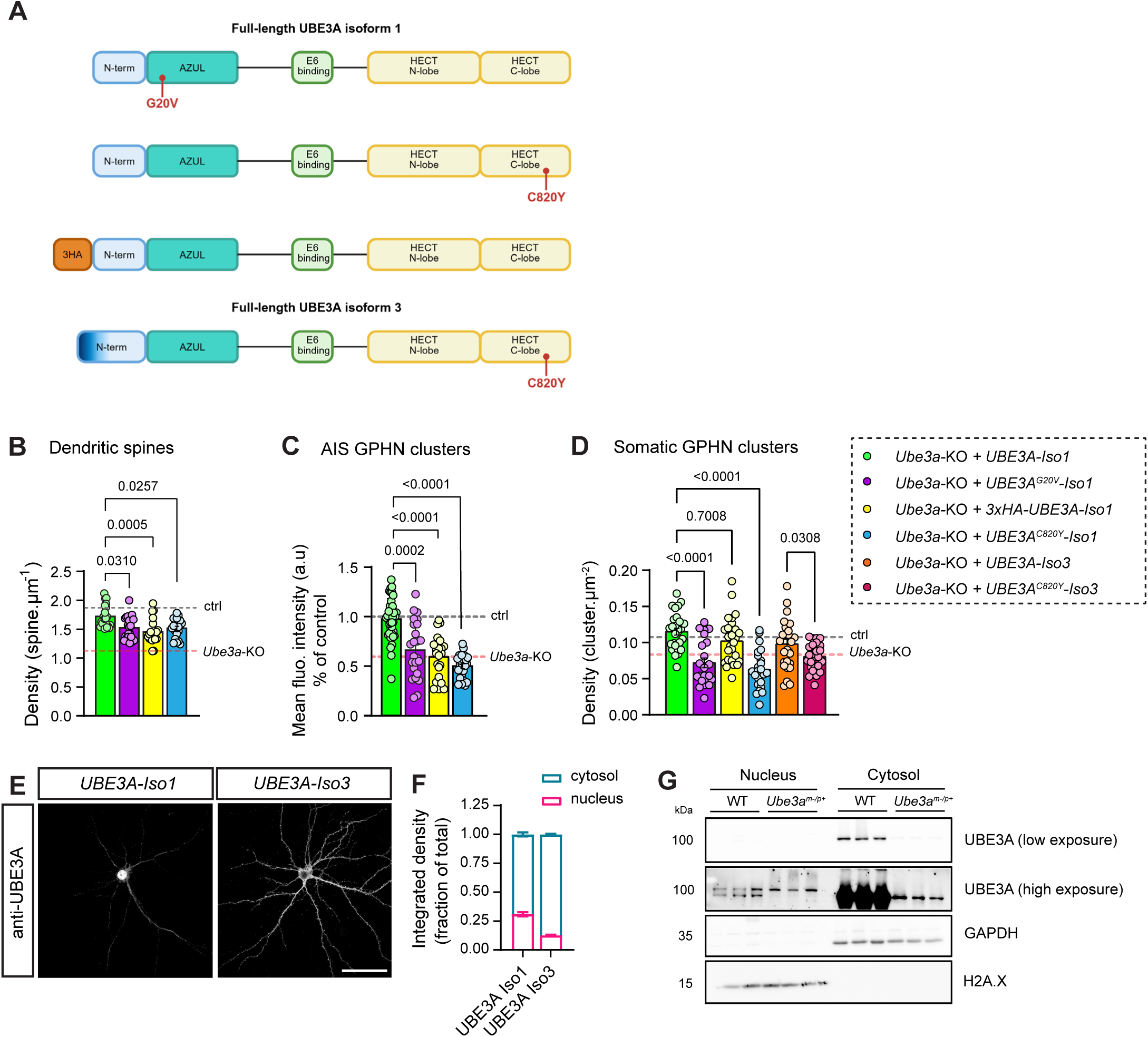
Molecular dissection of subcellular functions of UBE3A isoforms. **(A)** Schematic of UBE3A Isoform 1 (top) and Isoform 3 (bottom) functional domains and localization of the indicated mutations. N-term, N-terminal domain; AZUL, Amino-terminal Zn-finger of Ube3a Ligase domain; HECT, Homologous E6-AP Carboxy Terminus domain. Created in https://BioRender.com. **(B-D)** Quantification of dendritic spine density (B), normalized fluorescence intensity of AIS GPHN clusters (C) and density of perisomatic GPHN clusters (D) in CPNs after in utero replacement of endogenous *Ube3a* with UBE3A-Iso1 or Iso3 and the indicated mutants in juvenile mice. Dotted lines indicate control (grey) and *Ube3a*-KO (pink) values. Dendritic spines: n*_UBE3A-Iso1_* = 20 (4), n*_UBE3A G20V-Iso1_* = 21 (3), n*_UBE3A_ ^C820Y^_-Iso1_* = 21 (5); AIS GPHN clusters: n*_UBE3A-Iso1_* = 28 (3), n_UBE3A G20V-Iso1_ = 24 (4), n_UBE3A C820Y-Iso1_ = 24 (3); perisomatic GPHN cluster density: n_UBE3A-Iso1_ = 28 (3); n*_UBE3A G20V-Iso1_* = 24 (4); n*_UBE3A C820Y-Iso1_* = 24 (3); n*_UBE3A-Iso3_* = 26 (3); n*_UBE3A C820Y-Iso3_* = 25 (3). **(E-F)** Representative images (E) and quantification (F) of subcellular distribution and abundance of UBE3A in nucleus and cytosol in primary cortical neurons expressing UBE3A-Iso1 or UBE3A-Iso3 at DIV18. Scale bar: 50µm. n*_UBE3A-Iso1_* = 22 (3); n*_UBE3A-Iso3_* = 31 (3). **(G)** Representative western blot of UBE3A abundance in nuclear and cytosolic compartment upon subcellular fractionation of cortices from juvenile WT or *Ube3a^m−/p+^* mice. GAPDH and H2A.X are used as loading control and to test efficient enrichment of cytosolic and nuclear proteins, respectively, upon subcellular fractionation. Statistics: bars indicate mean ± SEM. Numbers in parentheses indicate the number of animals. p values are indicated in the graphs. Welch and Brown-Forsythe one-way ANOVA or Kruskal-Wallis test.

### Molecular determinants of UBE3A-dependent regulation of synapse development

To investigate the molecular bases of UBE3A isoform-specific regulation of synaptogenesis, we first assessed the importance of UBE3A intracellular localization. The amino-terminal Zn-finger of Ube3a ligase (AZUL) domain of UBE3A interacts with PSMD4, a subunit of the 19S regulatory particle of the 26S proteasome and dictates UBE3A nuclear localization (Avagliano Trezza *et al*, 2019). Thus, we generated an N-terminally 3xHA-tagged variant of UBE3A-Iso1, which was selectively redistributed from the nucleus to the cytosol without any change in its catalytic activity (Fig 6A; Figure S4) (Zampeta *et al*, 2020). Next, we used IUE to selectively replace endogenous *Ube3a* with this variant throughout development. The expression of 3xHA-UBE3A-Iso1 failed to rescue the formation of dendritic spines and the maturation of AIS inhibitory synapses (Fig 6B-C), highlighting the importance of UBE3A nuclear distribution for these subtypes of synaptic connections. Instead, CPNs expressing 3xHA-UBE3A-Iso1 displayed a density of perisomatic inhibitory synapses similar to that of control cells and neurons expressing WT UBE3A-Iso1 (Fig 6D). Together with our previous observation that UBE3A-Iso3 could restore GPHN cluster density in neuronal somata (Fig 5G-H), these results suggested that the presence of UBE3A in the cytosol is a critical determinant for the assembly of perisomatic inhibitory synapses.

To further interrogate the principles underlying UBE3A isoform-specific roles, we took advantage of previously characterized UBE3A mutants that were identified in AS patients, which display either an alteration of the subcellular localization of UBE3A (G20V mutant) or an impaired catalytic activity (C820Y mutant) (Fig 6A). Similar to the N-terminally tagged 3xHA-UBE3A-Iso1, the G20V mutation mislocalizes UBE3A-Iso1 from the nucleus to the cytosol (Figure S4) (Avagliano Trezza *et al*, 2019; Bossuyt *et al*, 2021; Yi *et al*, 2015). Instead, the C820Y mutation in the C-terminal HECT domain impaired UBE3A catalytic activity without impacting its subcellular distribution (Figure S4) (Bossuyt *et al*, 2021). We first dissected the nuclear function of UBE3A-Iso1. In juvenile mice, we found that the G20V mutant could not rescue dendritic spine density and the fluorescent intensity of GPHN clusters in the AIS, which remained significantly lower than in neurons expressing WT UBE3A-iso1 (Fig 6B-C), further substantiating the importance of UBE3A-Iso1 localization in the nucleus to ensure proper formation of excitatory synapses and maturation of AIS inhibitory synapses. We then analyzed perisomatic inhibitory synapses. Unexpectedly, the replacement of *Ube3a* with the cytosolically redistributed G20V UBE3A-Iso1 did not restore the density of somatic GPHN clusters, an observation that is seemingly at odds with results obtained using 3xHA-UBE3A Iso1 (Fig 6D). However, one possible explanation relies on the reported reduced protein stability of the G20V mutant (Zampeta *et al*, 2020; Sadhwani *et al*, 2018). To test this possibility, HEK cells were transiently transfected with WT or G20V UBE3A-Iso1 and subjected to a cycloheximide (CHX) chase assay for 6 hours. Protein levels of G20V UBE3A-Iso1 were strongly reduced compared to WT UBE3A-Iso1 before and after CHX exposure, thus indicating reduced expression levels of this mutant that might be primarily ascribed to impaired protein synthesis (Figure S4) (Bossuyt *et al*, 2021).

Finally, we tested whether UBE3A requires its catalytic activity to regulate synapses. By expressing the C820Y mutant, we demonstrated that both excitatory and inhibitory synapse development depends on UBE3A-mediated protein ubiquitination. Indeed, the expression of this ligase-dead mutant of UBE3A-Iso1 failed to restore the number of dendritic spines and the recruitment of GPHN molecules at AIS synapses (Fig 6B-C). Moreover, CPNs expressing either of the two C820Y mutant isoforms displayed impaired formation of perisomatic GPHN clusters (Fig 6D).

Our data showed that synaptic development depends on UBE3A functions in distinct subcellular compartments. In the nucleus, UBE3A-dependent ubiquitination regulates the assembly of excitatory connectivity and the maturation of AIS inhibitory synapses, while its cytosolic function mediated by either UBE3A-Iso1 or Iso3 is critical to ensure inhibitory synaptogenesis in the somatic region of CPNs. This last observation implied that UBE3A-Iso1 should partially reside in the cytosol, despite its reported nuclear enrichment (Avagliano Trezza *et al*, 2019; Zampeta *et al*, 2020; Sirois *et al*, 2020; Burette *et al*, 2017, 2018). To further analyze isoform-specific subcellular distribution of UBE3A, we parallelly employed imaging and biochemical approaches using cortical neurons and the most used AS mouse model (*Ube3a^m−/p+^*, heterozygous for *Ube3a*, maternal transmission) (Jiang *et al*, 1998). The latter was required to assess UBE3A localization in vivo using biochemical assays, which instead cannot be used with sparsely electroporated brains. First, we exogenously expressed individual UBE3A isoforms in dissociated cortical neurons (Fig 6E). To avoid background signal originating from endogenous *Ube3a*, it was knocked out using CRISPR/Cas9 genome editing (Figure S1). Using immunostaining, we then measured the integrated density of UBE3A fluorescent signal to assess the total abundance of UBE3A-Iso1 or Iso3 in the nucleus and cytosol. Although UBE3A-Iso1 was concentrated in the nucleus, ∼70% of UBE3A-Iso1 molecules were localized in the cytosol (Fig 6E-F). Instead, UBE3A-Iso3 was almost exclusively cytosolic, with more than 90% of molecules in the cytosol (Fig 6E-F). It is worth mentioning that the abundance of cytosolic UBE3A was measured excluding UBE3A signal from a region of interest (ROI) that was delineated using DAPI as reference. Vice versa, nuclear UBE3A was assessed quantifying its fluorescence in the same ROI. Thus, part of the signal defined as nuclear may instead come from the cytosol above and underneath the nucleus, potentially leading to underestimate and overestimate cytosolic and nuclear UBE3A localization, respectively. Next, we carried out subcellular fractionation and western blot analysis on cortices obtained from wild-type (WT) or *Ube3a^m−/p+^* mice. Considering that UBE3A-Iso1 constitutes ∼80% of total UBE3A (Sirois *et al*, 2020; Judson *et al*, 2021), this assay should allow to mainly infer on the localization of this isoform. Consistent with imaging data obtained from cultured cortical neurons, UBE3A expression in the cytosol largely exceeded that in the nucleus, where UBE3A was barely detectable without saturating UBE3A signal in the cytosolic fraction (Fig 6G). Altogether, our data indicated that in juvenile mice the number of UBE3A molecules is lower in the nucleus, where UBE3A is however concentrated and is crucial for excitatory synapses and AIS inhibitory synapses, and higher in the cytosol, where it regulates the formation of perisomatic inhibitory synapses.

### Impairments of synaptic development are recapitulated in *Ube3a^m−/p+^* mice

Sparse IUE is a method of choice to dissect intrinsic mechanisms of synaptic development in vivo (Fossati *et al*, 2022, 2016, 2019; Assendorp *et al*, 2024). Using this methodology, our results demonstrated that UBE3A controls the assembly and maturation of excitatory and inhibitory connectivity through cell-autonomous mechanisms. However, a limitation of IUE is the generation of mosaicism; in the case of our study, only a minority of cells suffered *Ube3a* deletion, a situation that is rarely encountered in AS patients, where *UBE3A* is lost in all neuronal cells (Nazlican *et al*, 2004). To test the translational relevance of our findings, we explored whether the observed synaptic phenotypes are also present in *Ube3a^m−/p+^* mice, which better mimic the genetic conditions of AS individuals (Jiang *et al*, 1998). *Ube3a^m−/p+^*animals were electroporated at E15.5 to express a fluorescent filler and FingR.GPHN, allowing to identify electroporated neurons, and to monitor dendritic spines and inhibitory synapses, respectively (Fig 7A). In line with data obtained from CRISPR/Cas9 electroporated animals, we found that ubiquitous *Ube3a* deletion affected the density of dendritic spines in layer 2/3 CPNs of the somato-sensory cortex of juvenile mice, which is reduced to 82% ± 3% compared to neurons from WT mice (Fig 7B-C). In *Ube3a^m−/p+^* animals, we also observed an impairment of the maturation of AIS inhibitory synapses (Figure 7D-E). The density of perisomatic inhibitory synapses also displayed a tendency towards a decrease in the absence of UBE3A, although it did not reach statistical significance (Fig 7F-G). Altogether, these data recapitulated the neuronal and synaptic phenotypes we identified using IUE and CRISPR/Cas9 editing to delete *Ube3a* in sparse CPNs, demonstrating the pathophysiological relevance of these synaptic alterations in AS.

**Fig 7.**
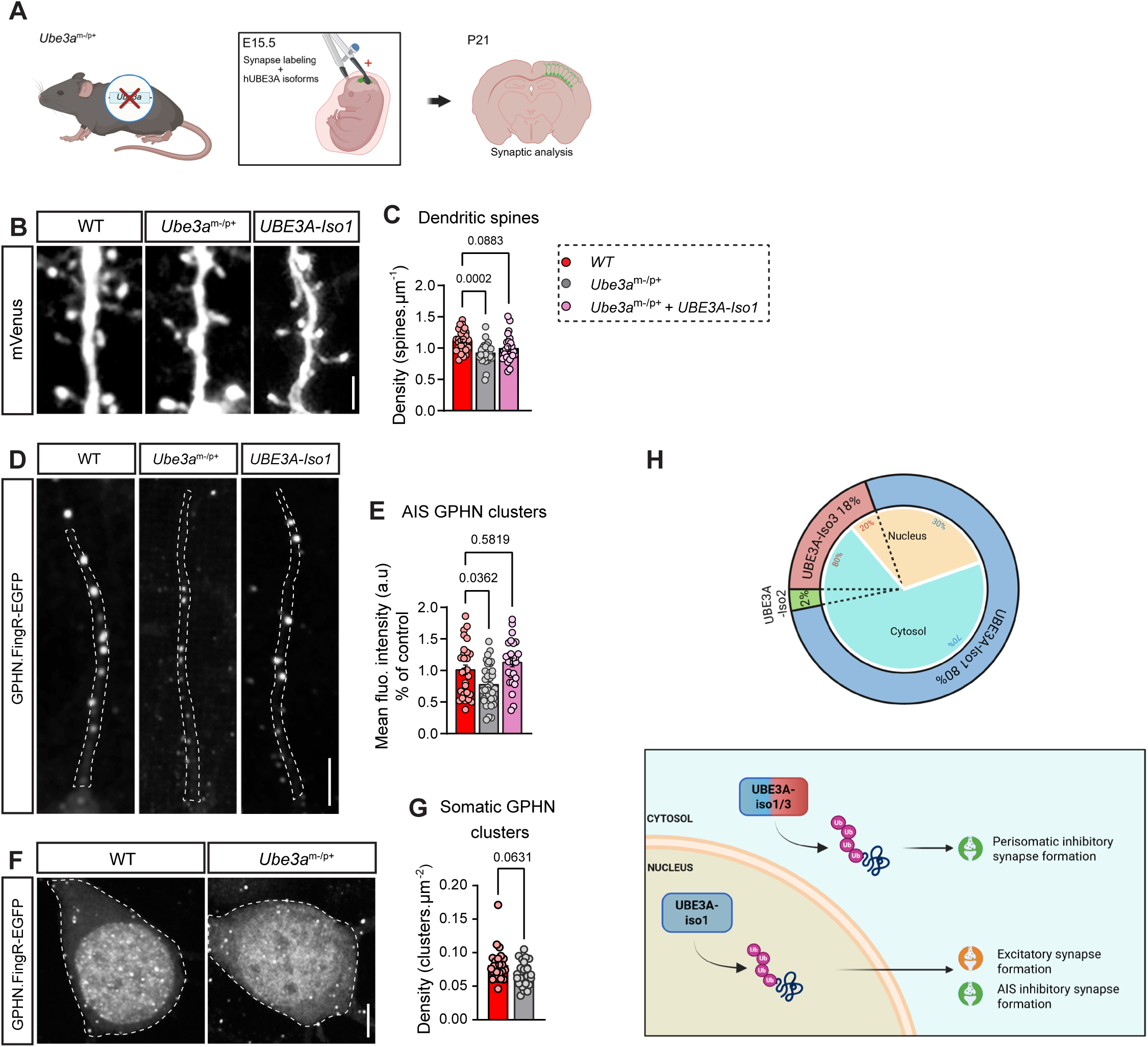
UBE3A-dependent alteration of synaptic development is driven by cell-autonomous mechanisms. **(A)** Schematic of sparse labeling of layer 2/3 CPNs after IUE in *Ube3a^m−/p+^*mice. Created in https://BioRender.com. **(B)** Segments of dendrites illustrating spines in juvenile WT and *Ube3a^m−/p+^* mice or upon expression of UBE3A-Iso1. Scale bar: 2 µm. **(C)** Quantification of spine density in conditions shown in(B). n_WT_ = 29 (3); n*_Ube3a m−/p+_* = 29 (5); n*_UBE3A-Iso1_* = 25 (3). **(D)** Representative images of AIS inhibitory synapses in neurons expressing GPHN.FingR-EGFP obtained from juvenile WT and *Ube3a^m−/p+^* mice or upon reinstatement of UBE3A-Iso1 using IUE. Dashed lines define the contours of AnkG fluorescence. Scale bar: 5 µm. **(E)** Quantification of normalized fluorescence intensity of AIS GPHN clusters in conditions shown in (D). n_WT_ = 32 (3); n*_Ube3a m−/p+_* = 37 (5); n*_UBE3A-Iso1_* = 27 (3). **(F)** Representative images of perisomatic inhibitory synapses in juvenile WT and *Ube3a^m−/p+^*mice. Dashed lines define the contours of DsRed fluorescence that was used to identify electroporated neurons and visualize neuronal morphology. Scale bar: 5 µm. **(G)** Quantification of perisomatic GPHN cluster density in conditions shown in (F). n_WT_ = 24 (3); n*_Ube3a m−/p+_* = 25 (5). **(H)** Left: schematic illustrating the expression (outer ring) and localization (central circle) of individual UBE3A isoforms. Right: schematic showing unique subcellular functions of UBE3A isoforms in the regulation of excitatory and inhibitory synaptic development (see main text for details). Created in https://BioRender.com. Statistics: bars indicate mean ± SEM. Numbers in parentheses indicate the number of animals. p values are indicated in the graphs. Welch and Brown-Forsythe one-way ANOVA or Kruskal-Wallis test.

If UBE3A governs synaptic development through cell-autonomous mechanisms, its reinstatement in sparse CPNs over a ubiquitous *Ube3a*-null background should be sufficient to restore excitatory synapse formation and AIS inhibitory synapse maturation. To test this possibility, we used IUE of *Ube3a^m−/p+^*animals and selectively expressed *UBE3A-Iso1* (Fig 7A). Strikingly, our results closely followed those observed in IUE-based *Ube3a*-KO mosaic cortices. Electroporation of *UBE3A-Iso1* enabled the rescue of dendritic spine density (Fig 7B-C) and fluorescence intensity of GPHN clusters located in the AIS (Fig 7D-E) to values similar to those observed in CPNs of WT animals. These experiments demonstrated that alterations of multiple developmental aspects of excitatory and inhibitory synapses onto layer 2/3 CPNs are consistently present across different mouse models of AS and further support the notion that they are governed by cell-intrinsic mechanisms.

## DISCUSSION

In the present study, we show that UBE3A is critical to the development and specification of excitatory and inhibitory connectivity in upper layers CPNs. We dissect the functional relevance of UBE3A isoforms that are generated by alternative splicing and are localized in distinct subcellular compartments, namely the nucleus and cytosol (Avagliano Trezza *et al*, 2019; Bossuyt *et al*, 2021; Sirois *et al*, 2020; van Esbroeck *et al*, 2025). We demonstrate that UBE3A intracellular compartmentalization, other than its molecular diversity, is crucial to regulate the development of distinct subtypes of synaptic connections, highlighting potential mechanisms underlying UBE3A function and AS pathophysiology.

### Synapse subtype-specific function of UBE3A

There is broad consensus that *UBE3A* genetic defects impair synaptic connectivity across neuronal networks (reviewed in (Biagioni *et al*, 2024)). The results presented here indicate that the formation of excitatory synapses onto layer 2/3 CPNs is significantly altered by the loss of *Ube3a*, ultimately affecting glutamatergic transmission at juvenile stages. This observation is in line with previous studies reporting decreased dendritic spine density in CPNs of the visual cortex in postmortem human samples (Jay *et al*, 1991) and various brain regions of AS mouse models (Sato & Stryker, 2010; Dindot *et al*, 2007; Wallace *et al*, 2012; Rotaru *et al*, 2023; Kim *et al*, 2016). However, increased excitatory transmission was described in layer 5 CPNs of the PFC and hippocampus (Rotaru *et al*, 2018; Kaphzan *et al*, 2011). These seemingly at odds observations might rely on different animal models that are used, and brain areas and developmental stages that are considered. The defective dendritic spine density at juvenile stages observed here indicates that this phenotype emerges during early postnatal neurodevelopment and may represent a premature cellular sign of AS. Interestingly, recent data assessing dorsomedial striatum developmental trajectories reported a stalled maturation that appears at the same developmental stage that we analyzed (Rotaru *et al*, 2023).

Besides impacting glutamatergic synapses, we show that *UBE3A* defects also alter the formation of inhibitory contacts, impairing the maturation of AIS synapse and the formation of perisomatic connections. Dysfunction of inhibitory connectivity is widely implicated in neurodevelopmental disorders, including Rett syndrome, which shares several behavioral phenotypes with AS (Ip *et al*, 2018). In AS, defective GABAergic transmission may produce enhancement of neuronal excitation in the cortex, leading to increased susceptibility to seizures (Rotaru *et al*, 2018; Wallace *et al*, 2012, 2017; Judson *et al*, 2016). In the present study, we demonstrate that *Ube3a* loss alters the development of specific subtypes of inhibitory synapses onto layer 2/3 CPNs, namely those located in neuronal somata and AIS involving presynaptic PV+ basket cells and PV+ chandelier cells, respectively (Tremblay *et al*, 2016). Among the diverse subtypes of inhibitory connectivity in the cortex, perisomatic and AIS synapses are closely located to the site of action potential generation and key determinants of CPN output, possibly contributing to hyperexcitable phenotypes despite the presence of fewer dendritic spines; this is confirmed by the increased E/I ratio in *Ube3a*-KO CPNs that we report in this study, and in line with previous observations in the visual cortex of adult AS mice (Wallace *et al*, 2012). The defective maturation of AIS inhibitory synapses are also accompanied by molecular and structural alterations of the AIS, possibly participating to enhanced neuronal excitability (Kaphzan *et al*, 2011; Wang *et al*, 2018). In this complex scenario, we cannot rule out primary versus secondary (homeostatic) changes. Longitudinal studies of developmental trajectories of synapses will clarify which synapse subtypes are directly targeted by *UBE3A* mutations at early postnatal stages or whether they are the consequence of compensatory mechanisms. The present study demonstrates that synaptic regulation by UBE3A occurs through cell-autonomous mechanisms. *Ube3a* inactivation in sparse layer 2/3 CPNs led to fewer functional synapses (excitatory contacts and perisomatic inhibitory synapses) and less mature AIS inhibitory synapses, likely due to postsynaptic effects. Although we cannot exclude that cell extrinsic mechanisms might additionally contribute to synaptic alterations in AS (Wallace *et al*, 2012), the recovery of synapse development in *Ube3a^m−/p+^* mice upon UBE3A reinstatement in single CPNs provide clear evidence that cell intrinsic mechanisms predominantly drive synaptic deficits in AS. Earlier studies reported an abnormal accumulation of clathrin-coated vesicles in excitatory and inhibitory terminals in the visual cortex of adult *Ube3a^m−/p+^* mice (Wallace *et al*, 2012; Judson *et al*, 2016). In Drosophila neurons, presynaptic UBE3A activity regulates synapse elimination through down-regulation of BMP signaling (Furusawa *et al*, 2023). These pre- and postsynaptic functions are not mutually exclusive and, instead, may synergistically operate to worsen synaptic function in AS. The temporal aspect is again of crucial importance and requires further investigation to determine primary versus secondary effects. For instance, defective synaptic vesicle cycling appears in adulthood (Wallace *et al*, 2012), while our observations are made in juvenile mice, raising the possibility that post-synaptic alterations might anticipate pre-synaptic vesicle impairment.

Using IUE to manipulate UBE3A expression and function at single-cell level, we limited our analysis to layer 2/3 CPNs, an approach that was crucial to isolate cell-autonomous mechanisms governing synaptic development in a cell-type specific fashion. Nonetheless, UBE3A is ubiquitously expressed in both excitatory and inhibitory neurons of both human and mouse brains (Judson *et al*, 2014; Burette *et al*, 2017). As aforementioned, earlier studies reported distinct functional alterations depending on cell types, cortical layers and areas that were analyzed (Rotaru *et al*, 2018; Wallace *et al*, 2012; Judson *et al*, 2016; Kaphzan *et al*, 2011; Rotaru *et al*, 2023; Dindot *et al*, 2007; Kim *et al*, 2016). Therefore, it is possible that UBE3A critically regulates the development and function of distinct synapse subtypes in a neuronal type-specific manner throughout different brain regions, whose dysfunction is associated with specific clinical impairments (Biagioni *et al*, 2024). Additional studies aiming to rigorously analyze these spatial aspects at multiscale will ultimately provide insights into subcellular and cellular bases of behavioral deficits in AS.

### The contribution of nuclear and cytosolic UBE3A to synapse development

Several immunohistochemical studies assessed the localization of UBE3A isoforms in various cellular systems, including immortalized cell lines, primary cultures of mouse neurons and IPSC-derived human neurons (Avagliano Trezza *et al*, 2019; Bossuyt *et al*, 2021; Sirois *et al*, 2020; van Esbroeck *et al*, 2025; Burette *et al*, 2017; Judson *et al*, 2021). Yet, the functional relevance of *UBE3A* molecular diversity and its subcellular localization to disease pathophysiology was poorly understood. Replacing endogenous *Ube3a* with individual isoforms, we show that UBE3A in the nucleus regulates the majority of synaptic alterations, namely the development of excitatory synapses and AIS inhibitory synapses. Accordingly, the reinstatement of UBE3A-Iso3 or mutants of UBE3A-Iso1 mislocalizing in the cytoplasm fail to restore either of these two phenotypes. The importance of nuclear UBE3A to AS pathophysiology was first suggested by the recent generation and characterization of isoform-specific KO mice (Avagliano Trezza *et al*, 2019), and then further highlighted following the identification of pathological mutations interfering with UBE3A localization in the nucleus without or only partially affecting its catalytic activity in AS individuals (Bossuyt *et al*, 2021). In line with this, a role for UBE3A in transcriptional regulation was described, but it was unclear whether this function depended on its ubiquitin ligase activity (Nawaz *et al*, 1999; Khan *et al*, 2006; Reid *et al*, 2003; Krishnan *et al*, 2017). Although in this study we do not directly assess UBE3A-dependent control of transcription, we demonstrate that nuclear UBE3A critically regulates the assembly of synaptic connectivity in a ubiquitin ligase-dependent fashion.

Employing variants of UBE3A, we also show that UBE3A in the cytosol contributes to the development of inhibitory connectivity, regulating the formation of perisomatic GPHN clusters. Remarkably, this is in line with other evidence suggesting the additional contribution of cytosolic UBE3A to neuronal development and AS deficits. In particular, the identification of two AS individuals carrying a point mutation selectively abrogating the start codon of nuclear UBE3A-Iso1 and displaying milder phenotypes suggested that cytosolic UBE3A-Iso2 and Iso3 are sufficient to attenuate disease severity (Sadhwani *et al*, 2018). Moreover, AS mice showed optimal rescue of behavioral deficits using a dual-isoform reinstatement strategy (Judson *et al*, 2021), and the selective expression of cytosolic UBE3A in *Ube3a*-KD hippocampal neurons prevented impairment of dendritogenesis (Miao *et al*, 2013). Very recently, a behavioral study using an animal model overexpressing cytosolic UBE3A (*mUbe3a-*Iso2) in an *Ube3a^m−/p+^* background was reported (Krzeski *et al*, 2026). Interestingly, the overexpression of cytosolic UBE3A could rescue a number of behavioral deficits that were assessed, but not those related to seizure/epileptogenesis. Although some differences that likely rely in the employed animal models and isoform-specific expression levels, these results are in line with our study indicating the presence of partially overlapping and partially isoform-specific functions of UBE3A. At present, it remains unclear why the G20V mutant does not behave as the 3xHA-UBE3A-Iso1 variant and fails to restore the proper formation of perisomatic inhibitory synapses. However, we show that the G20V mutation affects UBE3A protein expression, and others reported enhanced degradation (Sadhwani *et al*, 2018; Zampeta *et al*, 2020). UBE3A might not be sufficiently concentrated in the cytosol to carry out its function. Altogether, our data shed light on isoform-specific roles of UBE3A in synaptic development, potentially contributing to distinct circuit and behavioral alterations in AS.

### Subcellular distribution of UBE3A isoforms

Unexpectedly, this work indicates that the subcellular localization of UBE3A isoforms is less clear-cut than previously reported. The ability of both UBE3A-Iso1 and Iso3 to rescue the formation of perisomatic inhibitory synapses prompted us to inspect isoform-specific subcellular localization of UBE3A. Our imaging and biochemical data demonstrate that ∼70% of UBE3A-Iso1 molecules are distributed in the cytosol, where they are functionally interchangeable with UBE3A-Iso3 to mediate the recruitment of GPHN molecules in nascent perisomatic inhibitory synapses. As further confirmation, IUE of HA-tagged UBE3A-iso1, which is redirected in the cytoplasm, enables to restore the formation of perisomatic inhibitory connectivity. These results are seemingly at odds with previous work reporting that the shorter UBE3A-iso1 (or *mUbe3a-Iso3*) is predominantly enriched in the nucleus in mature neurons, while long hUBE3A-iso2 and 3 (or *mUbe3a-Iso2*) reside in the cytosol (Avagliano Trezza *et al*, 2019; Bossuyt *et al*, 2021; Judson *et al*, 2021; van Esbroeck *et al*, 2025). These apparent discrepancies may be explained by the relative nuclear and cytosolic volumes. In mature neurons, the nucleus occupies less than ∼10% of the total cell volume (Kandel, 2021), thus explaining why UBE3A is concentrated in the nucleus, although only a minor fraction of molecules resides there. Notably, UBE3A-Iso1 and Iso3 represent ∼80% and ∼18% of total UBE3A in neuronal cells, respectively (Judson *et al*, 2021). Moreover, our biochemical and imaging data indicate that ∼70% of UBE3A-Iso1 and more than 90% of UBE3A-Iso3 reside in the cytosol. Thus, UBE3A-Iso1 results the most concentrated and prevalent isoform in this compartment (Figure 7H). Consistent with these results, localization of UBE3A-Iso1 in the cytoplasm was already observed using subcellular fractionation of isoform-specific KO human neurons derived from induced pluripotent stem cells (Sirois *et al*, 2020).

These observations also raise fundamental questions about the mechanisms underlying the functional roles of UBE3A isoforms. Do they possess specific affinities for distinct substrates or interact with different proteins because of their uneven subcellular compartmentalization? Our in utero gene replacement strategy introducing targeted mutations points toward the second hypothesis. The ability of 3xHA-UBE3A-Iso1, which is excluded from the nucleus and redirected to the cytosol, to rescue perisomatic inhibitory synapses provides a strong indication that UBE3A isoforms likely interact with the same targets when they are distributed in the same compartment. This is also consistent with a very recent study showing that overexpression of cytoplasmic UBE3A could rescue a number of behavioral deficits in mice (Krzeski *et al*, 2026). In a translational perspective, this knowledge is critical for potential therapies based on UBE3A gene transfer, which must consider UBE3A subcellular localization as major determinant of UBE3A function. Future studies employing isoform-specific transgenic animals combined with proteomics across neurodevelopment will help to clarify this aspect and identify substrates of individual isoforms.

In conclusion, the present work uncovers unique roles of UBE3A isoforms in specific subcellular compartments, which are crucial for the assembly of neuronal circuits, ultimately unlocking new pathways to understand disease mechanisms and inform targeted therapeutic strategies for AS.

## METHODS

### Animals

All animals were handled according to EU regulations and were approved by the Italian Ministry of Health (Authorization numbers 596/2019-PR and 750/2021-PR). In utero electroporations were performed on pregnant CD1 females (Charles River Laboratory) or heterozygous C57BL/6J *Ube3a^p−/m+^*dams (giving birth to either WT or *Ube3a^m−/p+^* mice, The Jackson Laboratory) (Jiang *et al*, 1998) at E14.5-15.5. Primary cultures were prepared from timed pregnant CD1 mice at E18.5 (Charles River Laboratory). Juveniles correspond to mice between P21 and P23, with the day of birth defined as P0. Mice were maintained in a 12 hr light/dark cycle with unlimited access to food and water. In all experiments, data were collected without considering the sex of the animals.

### DNA constructs for mammalian gene expression

All plasmids for gene expression through IUE had a pCAG backbone driving protein expression under the CAG promoter. For pCAG UBE3A-EGFP isoforms, human *UBE3A-Iso3* cDNA was obtained from Horizon (Clone id: 3160225; GenBank: BC002582.2), subcloned by PCR and inserted into pCAG between KpnI and AgeI; pCAG UBE3A-EGFP Isoform 1 was subcloned by PCR from UBE3A isoform 3 to remove the first N-terminal 20aa of isoform 3 and inserted between HindIII and AgeI. Untagged pCAG UBE3A-Iso1 and Iso3 were subcloned by PCR to remove the C-terminal EGFP tag, add a stop codon and inserted into pCAG between SacI and NotI. pCAG UBE3A-T2A-tdTomato isoforms were generated by inserting T2A-tdTomato into pCAG UBE3A-Iso1 and Iso3 downstream UBE3A using InFusion cloning (Takarabio). For pCAG 3xHA-UBE3A, a DNA cassette containing a start codon and 3xHA tags was inserted into pCAG UBE3A-Iso1 between HindIII and SacI. Mutants of UBE3A isoforms (G20V and C820Y) were obtained by site-directed mutagenesis (Agilent). pCAG PSD95.FingR-EGFP-CCR5TC and pCAG GPHN.FingR-EGFP-CCR5TC were a gift from Don Arnold (Addgene plasmids #46295 and #46295) (Gross *et al*, 2013). All constructs were verified by DNA sequencing.

### Generation and validation of plasmids for CRISPR/Cas9- and shRNA-mediated gene inactivation

To knock out *Ube3a* with CRISPR, we used an engineered spCas9 with enhanced specificity (espCas9 (1.1)), gift from Feng Zhang, Addgene plasmid #71814) (Slaymaker *et al*, 2016). gRNAs were designed using the prediction software: https://portals.broadinstitute.org/gpp/public/analysis-tools/sgrna-design. *Ube3a* was knocked-out using a bicistronic plasmid containing one gRNA under the control of the U6 promoter together with the Flag-tagged espCas9 (1.1) under the CAG promoter. Control CRISPR consisted in the plasmid without any gRNA. A DNA cassette encoding the gRNA against *ube3a* was inserted between BbsI sites. The gRNA targets the following sequence of exon 6 of *ube3a*: 5’- GATCCACTAGAAACCGAACT-3’. Importantly, this gRNA targeted the mouse *ube3a* gene only, and not the human paralog, which was used in rescue experiments using the in utero gene replacement strategy. CRISPR/Cas9-mediated deletion of *ube3a* was first validated *in vitro* using primary cultures of cortical neurons. Briefly, dissociated neurons at days in vitro (DIV)7 were transfected with the espCas9 (1.1) either without any gRNA or with the gRNA against *Ube3a*. At DIV14, neurons were fixed, stained using an anti-Flag antibody to reveal Cas9 expressing cells and anti-UBE3A. For in vivo validation, pregnant CD1 females were co-electroporated in utero at E14.5-15.5 with control or gRNA *ube3a* espCas9 (1.1) plasmids and the pCAG DsRed to identify transfected neurons. At P21, mice were intracardially perfused. Brain slices obtained at the vibratome were then subjected to immunohistochemistry using an anti-UBE3A antibody. For in utero knockdown experiments with shRNAs, we used the previously described pH1SCV2 and pH1SCTdT2 vectors (Charrier *et al*, 2012; Fossati *et al*, 2016, 2019). An H1 promoter drives the expression of the shRNA and a CAG promoter that of myristoylated Venus (mVenus) or TdTomato, respectively. The vector pH1SCV2 was used for dendritic spine analysis, pH1SCTdT2 was co-expressed with PSD-95 and GPHN FingRs to analyze excitatory and inhibitory synapses. For shRNA validation on endogenous mouse *ube3a*, we used a lentiviral vector carrying the H1 promoter to drive shRNA expression and the synapsin promoter to drive EGFP expression (Fossati *et al*, 2019, 2016). The control shRNA (shControl) was 5’-ACACCTATAACAACGGTAG-3’(Fossati *et al*, 2019). The seed sequence for *ube3a* was 5’-GCCCAGACACAGAAAGGTTAC-3’. A DNA cassette containing the shRNA against *Ube3a* was inserted into the lentiviral vector between BamHI and HindIII sites. For the knockdown validation, primary cultures of cortical neurons were infected at DIV4 with lentiviruses carrying an shRNA against *Ube3a* or a control shRNA. Infected cultures were harvested at DIV18 and lysed in RIPA buffer under agitation for 1h at 4°C and further processed for western blot analysis. All constructs were verified by DNA sequencing.

### Lentivirus production and purification

48 h after transfection of HEK293T cells, the viral supernatant was collected and subsequently concentrated using Lenti-X Concentrator (Clontech), according to manufacturer protocol. After brief centrifugation of viral supernatant to remove cell debris, Lenti-X Concentrator was added in a 3:1 ratio and incubated for 4h at 4°C. Viral particles were then centrifuged at 1,500 x *g* for 45 min at 4°C and the pellet resuspended in sterile PBS, aliquoted and stored at −80°C. When indicated, cortical neurons were infected 4 days after plating with concentrated lentiviruses driving the expression of shRNA and EGFP.

### Primary cultures of cortical neurons

Primary cultures were performed as described previously with few modifications (Charrier *et al*, 2012). After dissection and dissociation of mouse cortices from E18.5 embryos, neurons were plated on glass coverslips or directly on dishes coated with poly-D-ornytine (80 mg/ml, Sigma) in MEM supplemented with sodium pyruvate, L-glutamine (2 mM) and 10% horse serum. Medium was changed 2 h after plating with Neurobasal supplemented with L-glutamine (2 mM), B27 (1X) and penicillin (2.5 units/ml) - streptomycin (2.5 mg/ml). Then, one third of the medium was changed every 5-6 days. Unless otherwise indicated, all products were from Life Technologies. Cells were maintained at 37°C in 5% CO_2_ until use.

### HEK cells

HEK293T (CRL-1573 from ATCC) cells were cultured according to suggested protocols. Briefly, cells were maintained in DMEM (Sigma) supplemented with 10% fetal bovine serum (VWR) and 1% Penicillin-Streptomycin (GIBCO) at 37°C, 5% CO_2_, and passaged by trypsin/EDTA digestion (Euroclone) upon reaching confluency. A maximum of 30 consecutive passages were carried out before thawing a new cell aliquot.

### Transfection

Transfection was performed using Lipofectamine 2000 (Invitrogen) for both neuronal cultures and HEK cells according to manufacturer protocol with few modifications for cortical neurons. Briefly, we collected conditioned medium from dissociated cortical neurons at DIV7-9 and replaced with fresh Neurobasal without any supplement. DNA and lipofectamine were diluted in Neurobasal in separate tubes, and then lipofectamine mix was added to DNA mix. After 15 min incubation at RT, DNA-lipofectamine mix was added to cortical neurons and incubated for 1h at 37°C, 5% CO_2_. Next, conditioned medium was put back to neurons until use at DIV14-18.

### Immunocytochemistry

At DIV17-18, primary culture of cortical neurons were fixed for 15 min at room temperature (RT) using 4% (w/v) paraformaldehyde (PFA) in PBS and incubated for 30 min in blocking buffer containing 0.3% Triton X-100 (Sigma) and 3% Bovine Serum Albumine (BSA from Sigma). Subsequently, cells were incubated for 1h in primary antibody, rinsed 3 x 5 min in 1x PBS, and incubated 45 min in secondary antibodies diluted in blocking buffer. After washing first in 0.01% Triton X-100 in 1x PBS and then twice in 1x PBS, coverslips were rinsed in distilled H_2_O and mounted on slides with Vectashield or Mowiol (Sigma). For cell surface staining, dissociated neurons at DIV14-18 were incubated for 15 min in primary antibodies diluted in imaging medium (Sigma) at 37°C. After rinsing in imaging medium, neurons were fixed for 5 min at RT using 4% PFA in 1x PBS and further processed for standard immunocytochemistry. Primary antibodies were rabbit anti-α2 GABA_A_R (Synaptic systems, 1:500), anti-©2 GABA_A_R (Synaptic Systems, 1:500), mouse anti-PSD-95 (Neuromab, 1:1000), mouse anti-Gephyrin (Synaptic Systems Clone 7a, 1:800), mouse anti-FLAG (Cell Signaling Technologies, 1:800), mouse anti-UBE3A (Bethyl Laboratories,1:250) and guinea pig anti-ankyrin G (Synaptic Systems, 1:500). Secondary antibodies were Alexa-conjugated (Invitrogen) or Abberior STAR (Abberior) and diluted 1:500.

### In utero electroporation and slice preparation

IUE was carried out as follows (Fossati *et al*, 2022, 2019, 2016). Pregnant CD1 females at E14.5-15.5 (Charles River) were anesthetized with isoflurane (3.5% for induction and 2% with air flow at 1L/min for maintenance during surgery) and subcutaneously injected with 5 mg/kg Flunixin Meglumine for analgesia. The uterine horns were exposed after laparotomy. Electroporation was performed using a square wave electroporator (Nepagene) and tweezer-type platinum disc electrodes (5mm-diameter, Sonidel). The electroporation settings were: 4 pulses of 40 V for 50 ms with 500 ms interval. Endotoxin-free DNA was injected into one lateral ventricle of the mouse embryos using a glass pipette obtained pulling borosilicate microcapillaries with 1.5mm diameter (World Precision Instrument). The volume of injected DNA was adjusted depending on the experiments. Plasmids were used at the following concentrations: CRISPR/Cas9 knock out plasmids: 1 mg/ml; pCAG dsRed: 0.5 mg/ml, pCAG TdTomato: 0.5 mg/ml, pCAG EGFP: 0.5mg/ml, pCAG mVenus: 0.5mg/ml, GPHN.FingR-EGFP and PSD95.FingR-EGFP: 0.7 mg/ml; shRNA vectors: 1 mg/ml; pCAG plasmids driving the expression of UBE3A isoforms (Iso1 and Iso3) and variants (tagged, untagged, T2A-tdTomato, G20V, C820Y and T485A mutations): 1 mg/ml. Animals were sacrificed at the indicated age by intracardiac perfusion of 4% PFA in PBS (Electron Microscopy Sciences). Unless otherwise indicated, 100 μm coronal brain sections were obtained using a vibrating microtome (Leica VT1200S, Leica Microsystems). Sections were mounted on slides in Vectashield (Vector Laboratories).

### Immunohistochemistry

100μm- or 50μm-thick coronal sections obtained at the vibrating microtome were processed for immunohistochemistry as follows. Briefly, free-floating slices were incubated in 0.1% TritonX-100 and 4% normal goat serum (NGS) (Sigma-Aldrich) in PBS to permeabilize and block unspecific staining. Primary antibodies were incubated overnight (O/N) at 4°C and secondary antibodies for 3h at RT under gentle agitation. Both primary and secondary antibodies were diluted in 0.1% Triton X-100 and 2% NGS in PBS. After each incubation slices were extensively rinsed in 1x PBS. Slices were mounted on slides in either Vectashield (Vector Laboratories) for confocal microscopy or Mowiol (Sigma) for STED microscopy. Guinea pig anti-ankyrin G (Synaptic Systems, 1:500), mouse anti-VGAT (Synaptic Systems, 1:800), mouse anti-UBE3A (Sigma, 1:1,000) and mouse anti-GFP (Roche) were used as primary antibodies. All Alexa-conjugated secondary antibodies (Invitrogen) or Abberrior STAR (Abberior) were diluted 1:500.

### Confocal and wide-field imaging

Confocal images were acquired in 1024×1024 mode using either a Leica TCS SP8 or a Leica TCS SP5 scanning confocal microscopes controlled by the LAS AF or LASX software, respectively, and equipped with a laser unit including 405 nm, 488 nm, 555 nm and 633 nm lasers and hybrid detectors (Leica Microsystems). We used a 20x objective (NA 0.7) to acquire low magnification images and 40x (NA 1.25) or 63x (NA 1.4) objectives to acquire high magnification images for synaptic analysis. Z-stacks were acquired with 150 nm spacing and a zoom of x1.24. To assess UBE3A subcellular distribution in the nucleus and cytosol, wide-field images were acquired using a Leice Thunder microscope controlled by the LAS AF software and equipped with monochromatic light emitting diodes, including 395 nm, 475 nm, 511 nm, 555 nm, 575 nm and 635 nm wavelengths and a CMOS camera (Leica DFC9000 GT, Leica Microsystems).

Confocal images of brain slices were acquired in 1024×1024 mode using either a Leica TCS SP8 or a Leica TCS SP5 scanning confocal microscopes controlled by the LAS AF or LASX software, respectively, and equipped with a laser unit including 405 nm, 488 nm, 555 nm and 633 nm lasers and hybrid detectors (Leica Microsystems). We used a 10x or 20x objective (NA 0.40 and 0.80, respectively) to identify electroporated neurons and acquire low magnification images, and a 63x objective (NA 1.4) objective to acquire higher magnification images and z-stacks of neuronal dendrite, soma and AIS. For synaptic analysis, Z-stacks were acquired with 190 nm spacing and a zoom of x 1.5 or x 2.0. Images were blindly acquired and analyzed.

### STED microscopy

For STED microscopy *xyz* images were acquired with a Leica TCS SP8 STED3X confocal microscope using a 100x/1.40 oil objective and a pixel size of approximately 38×38×70nm (x-y-z) to achieve an appropriate sampling interval for deconvolution. Electroporated neurons were excited with 488nm-tuned White Light Laser (WLL) and emission collected from 495nm to 593nm with a HyD detector. Gated 660nm CW-STED depletion laser was applied to 488nm-tuned WLL. Images were subsequently deconvolved with Huygens Professional software (Scientific Volume Imaging, version 21.04).

### Western blot

SDS-PAGE was carried out using 10% gels (Biorad). Western blotting was performed using the following primary antibodies: mouse anti-HA (Cell Signaling Technologies, 1:2,000), rabbit anti-GFP (Life Technologies, 1:2,000), rabbit anti-RFP (Rockland Immunochemicals, 1:1,000), rabbit anti-GAPDH (Synaptic Systems, 1:1,000), mouse anti-UBE3A (Sigma, 1:1,000), mouse anti-β3 GABA_A_R (Neuromab, 1:1,000), mouse anti-GluA1 (Cell Signaling Technologies, 1:300), mouse anti-GluA2 (Synaptic Systems, 1:500), mouse anti-Transferrin Receptor (ThermoFisher, 1:1000) and guinea pig anti-ankyrin G (Synaptic Systems, 1:500). All HRP-conjugated secondary antibodies were used at 1:10,000 dilution (ThermoFisher). Protein visualization was performed by chemiluminescence using Supersignal PICO Super or FEMTO Maximum western blotting substrates (ThermoFisher) and Chemidoc (Biorad) imagers.

### Cell surface biotinylation

After infection with lentiviral particles to knock-down *Ube3a*, primary cultures of cortical neurons at DIV17 were washed 3 times in ice-cold PBS supplemented with 0.8 mM CaCl_2_ and 0.5 mM MgCl_2_ (PBS^2+^) and then incubated for 5 min at room temperature followed by further 10 min at 4°C with 0.5 mg/ml Sulfo-NHS-SS-Biotin (ThermoFisher Scientific) in PBS^2+^. After rinsing in ice-cold PBS^2+^, biotin was quenched in 50 mM glycine in PBS^2+^ for 10 min. Next, cells were scraped in NaCl-Tris buffer (150 mM NaCl, 50 mM TrisHCl pH 7.4) supplemented with protease inhibitory cocktail (Roche) and lysed in extraction buffer (150 mM NaCl, 50 mM TrisHCl, 1% Triton X-100, 10 mM EDTA, protease inhibitor cocktail) for 1h at 4°C. Biotinylated proteins were pulled down by incubating cell lysates with streptavidin magnetic beads (ThermoFisher Scientific) for 3h at 4°C. After rinsing 3 times in extraction buffer and twice in scraping buffer, beads were resuspended in gel loading buffer (Sigma) and bound proteins were eluted with boiling. Relative cell surface expression levels were analyzed by western blotting. Inputs correspond to 10% of the cell surface fraction.

### UBE3A stability assay

Protein stability of UBE3A variants was probed in HEK cells using cycloheximide as described in (Geerts-Haages *et al*, 2020). Briefly, HEK293T cells were transiently transfected to drive the expression of UBE3A-Iso1 variants. 48 h post-transfection, cells were directly harvested (t_0_) or treated with 50 μg/ml cycloheximide (Sigma) for 6 h (t_1_) before harvesting. Upon brief centrifugation (5 min, 1,600 x *g*), cell pellets were lysed in lysis buffer (50mM TrisHCl pH 7.4, 150mM NaCl, 1% Triton X-100, 2 mM EDTA pH 8 and protease inhibitor cocktail). Protein lysates (15 μg) were analyzed by SDS-PAGE and western blotting.

### Ubiquitination assay

UBE3A auto-ubiquitination capacity of different UBE3A variants was evaluated using a cell-based ubiquitination assay. Briefly, HEK293T cells were co-transfected with a pCAG plasmid driving the expression of UBE3A-Iso1 variants and a second plasmid for EGFP-ubiquitin (Addgene, #187910). Cells were grown for 48 h and treated with 30 µM MG-132 (Sigma) for 2 h. Cells were harvested and lysed in lysis buffer (50mM TrisHCl pH 7.4, 150 mM NaCl, 1% Triton X-100, 2 mM EDTA pH 8, 1x protease inhibitor cocktail) supplemented with 30 μM MG-132. Cell lysates (1mg of protein/sample) were immunoprecipitated with mouse anti-UBE3A antibody (Sigma) O/N at 4°C under rotation. Immunoprecipitates were incubated with Protein G magnetic beads (ThermoFisher; cat) and incubated for 3h at 4°C under rotation. The final complex was washed three times in wash buffer (50 mM TrisHCl pH 7.4, 200mM NaCl, 1% Triton X-100, 2 mM EDTA pH 8, 1x protease inhibitor cocktail), resuspended in sample buffer and processed for SDS-PAGE and immunoblot analysis.

### Subcellular fractionation

After dissection, brain tissues were placed in C Tubes (Miltenyi Biotec) and homogenized in hypotonic buffer (10 mM Hepes-KOH pH 7.9, 10 mM Tris-HCl pH 7.4, 10 mM KCl, 1.5 mM MgCl2, 0.5 mM DTT) supplemented with protease inhibitors (Roche) and 13.4 mM NEM (Sigma) using the Program “4C_nuclei” on GentleMACSTM Octo Dissociator (Miltenyi Biotec). The amount of hypotonic buffer is related to the weight of the tissue according to manufacturer instruction (Miltenyi Biotec, “Nuclei Extraction Buffer”). After homogenization, brains were added with 0,3 % (v/v) Igepal (Sigma) and gently pipetted up and down. Whole homogenates were filtered with 70 μm-cell strainers and washed with 500 μL of hypotonic buffer supplemented with 0,3% (v/v) Igepal. Samples were centrifuged for 5 minutes at 500 x *g* at 4°C and supernatant (S1, cytosolic fraction) was separated from the resulting pellet (P1, nuclear fraction). S1 supernatant was diluted 1:1 in lysis buffer (50 mM Tris-HCl pH 7.5, 150 mM NaCl, 5 mM MgCl2, 1% Igepal, 0.5% DOC, 0.1% SDS) supplemented with protease inhibitors and incubated under constant rotation for 1h at 4°C. Samples were then sonicated 3 times for 10 seconds (off time 30 sec) and centrifuged at 16,000 x *g* for 10 minutes at 4°C. Supernatants were collected and quantified with microBCA kit (Thermo Scientific) for subsequent western blot analysis. P1 pellets were resuspended in 1 mL of isotonic buffer (50 mM Tris-HCl pH 7.5, 150 mM NaCl, 5 mM MgCl2) supplemented with protease inhibitors and 13.4 mM NEM, filtered with 40 μm-cell strainers and washed with 500 μL of isotonic buffer. Suspensions were centrifuged at 10,000 x *g* for 7 minutes at 4°C and supernatants were removed. Pellets were further resuspended in 500 μL of lysis buffer and incubated under constant rotation for 1h at 4°C. Samples were heated at 55°C for 10 minutes and sonicated 3 times for 30 seconds (off time 30 sec). Finally, they were centrifuged at 16,000 x *g* for 10 minutes at 4°C and supernatants were quantified with microBCA kit and used for subsequent experiments.

### Electrophysiology

Electroporated mice at P21 were deeply anesthetized with isoflurane at 4% by inhalation and decapitated. Brains were removed and placed in an ice-cold cutting solution containing the following (in mM): 87 NaCl, 21 NaHCO3, 1.25 NaH2PO4, 7 MgCl2, 0.5 CaCl2, 2.5 KCl, 25 D-glucose, and 7 sucrose, equilibrated with 95% O2 and 5% CO_2_ (pH 7.4). Coronal slices (300 mm thick) were cut with a VT1000S vibratome (Leica Microsystems). Slices were incubated for at least 1 h at 37°C in artificial cerebrospinal fluid (ACSF) containing the following (in mM): 129 NaCl, 21 NaHCO3, 1.6 CaCl2, 3 KCl, 1.25 NaH2PO4, 1.8 MgSO4, and 10 D-glucose, aerated with 95% O2 and 5% CO2 (pH 7.4). During recordings, slices were continuously perfused with ACSF at a flow rate of 1.5 ml/min. Recordings were carried out on electroporated cells localized in layer 2/3 somato-sensory cortex, which were visually identified using an Olympus BX51WI upright microscope equipped with water-immersion 10x and 40x objectives and an infrared (IR) camera (XM10r, Olympus). Whole-cell patch-clamp recordings were performed using a Multiclamp 700B amplifier (Molecular Devices) at 30°C using an Automatic Heater Controller (Warner Instruments Model TC-344C Heater Controller). Low-resistance micropipettes (4–6 MΩ) were pulled from borosilicate glass pipettes. Signals were low-pass filtered at 2 kHz, sampled at 20 kHz, and analyzed using a Digidata 1550B system (Molecular Devices). Basal excitatory and inhibitory synaptic transmission was simultaneously recorded from the same cell in the presence of 1 mM tetrodotoxin (TTX). mEPSCs and mIPSCs were recorded at a holding potential of –70 mV and +20 mV, respectively. Recordings were considered reliable if the following criteria were met: *i)* series resistance was below 25 MΩ (typically between 10 and 12 MΩ); *ii)* series resistance did not change by more than 20% during the recording; and *iii)* holding current did not exceed –150 pA. Series resistance was not compensated. The internal pipette solution contained (in mM): 138 Cs-gluconate, 2 NaCl, 10 HEPES, 4 EGTA, 0.3 Tris-GTP, and 4 Mg-ATP (pH adjusted to 7.2).

### Image analysis of immunocytochemistry experiments

In immunocytochemistry experiments, the abundance of surface g2- and a2-GABA_A_R subunits at AIS synapses was analyzed manually. AIS was identified using AnkG labeling and its length was measured drawing a segment delineating AnkG-positive area. Regions Of Interest (ROIs) were manually drawn around synaptic clusters in each channel. Morphometric parameters were measured for γ2- and α2-GABA_A_R clusters colocalizing with GPHN puncta.

The subcellular distribution of UBE3A isoforms was assessed using Fiji (https://fiji.sc/). A user-defined intensity threshold was applied to tdTomato channel to define a ROI that delineates the entire neuronal morphology. Nuclear surface was similarly annotated using a user-defined threshold in Hoechst channel. Cytosolic surface was obtained subtracting nuclear surface from tdTomato-based ROI. Such defined surface profiles were used to measure the mean and integrated fluorescence intensity of individual UBE3A isoforms in nuclear and cytosolic compartments.

### Image analysis of brain slices

To assess the efficiency of CRISPR-based deletion of *Ube3a* in vivo, the mean fluorescence intensity of UBE3A was quantified using Fiji. ROIs were drawn to delineate contours of neuronal somata of electroporated cells based on cytosolic tdTomato fluorescence and applied to UBE3A signal. After background subtraction, UBE3A fluorescent signal was measured in each electroporated cell, and then averaged for each brain slice. Dendritic spines, PSD-95 and GPHN clusters were analyzed in different sets of neurons and neuronal subcompartments using Fiji. Spines were quantified based on mVenus fluorescence conveyed by the pH1SCV2 vector. GPHN and PSD-95 clusters were quantified based on EGFP fluorescence of GPHN and PSD-95 FingRs in electroporated neurons identified for the expression of soluble DsRed or TdTomato, driven by pCAG DsRed or bicistronic T2A-tdTomato vectors, respectively. All quantifications were performed in layer 2/3 of the somatosensory cortex, in sections of comparable rostro-caudal position. PSD-95 clusters and dendritic spines were quantified in the proximal part of oblique dendrites directly originating from the apical trunk. Only dendrites that were largely parallel to the plane of the slice were analyzed to limit underestimation of dendrite length, which was measured on *z* projections. One dendrite per cell was analyzed. The density of dendritic spines and PSD-95 clusters along dendrites of a minimal length of 60 μm was calculated as described (Fossati *et al*, 2016, 2019; Assendorp *et al*, 2024). GPHN clusters were quantified in distinct subcellular compartments of layer 2/3 CPNs, namely dendrites, soma and axon initial segment (AIS). GPHN clusters in the proximal part of oblique dendrites were assessed as previously described for dendritic spines and PSD-95 (Fossati *et al*, 2019; Assendorp *et al*, 2024; Fossati *et al*, 2016). GPHN clusters in the AIS were identified using AnkG immunostaining. The density of GPHN clusters was quantified by manually counting the number of GPHN puncta in the AIS. Since images were acquired with different confocal microscopes, the fluorescence intensity of GPHN FingRs was expressed as a fraction of the average intensity observed in control neurons (normalized fluorescent intensity). The position of GPHN clusters in AIS was measured as the percentage of clusters in the first half of AIS. For perisomatic inhibitory synapses, co-localizing GPHN and vGAT clusters on the surface of TdTomato positive neurons were manually counted. Imaris 9.0 software (Bitplane AG) was used to detect surfaces in TdTomato channel and generate a 3D mask of the soma. GPHN and vGAT density was calculated dividing the number of detected clusters by soma volume. Fiji software was used for morphometric analysis of STED images of dendritic spines. Segments corresponding to spine neck and head dimensions were manually drawn and measured. All images were blindly analyzed.

### WB analysis

Protein visualization was done by chemiluminescence using Chemidoc imager (Biorad). When needed, optical density was quantified using Fiji. The signal associated with the protein of interest was normalized to either GAPDH expression, which was used as housekeeping reference for total and cytosolic homogenates, RFP expression, used as control of transfection efficacy, or H2A.X expression for nuclear homogenates. A minimum of 3 replicates for each experiment was performed.

### Statistics

Statistical analyses were performed with Prism 9 and Prism 10 software (GraphPad). All experiments were carried out in at least 3 independent replicates. For IUE, data were obtained from at least 3-4 animals from 2-3 independent litters. For statistical analysis, normality of the distributions was assessed using the D’Agostino-Pearson normality test. Comparisons between two groups were made using either unpaired two-tailed Student’s t-tests, when normally distributed, or non-parametric Mann-Whitney test, in case of non-normal distribution. Welch and Brown-Forsythe one-way ANOVA followed by Tukey’s post test was used to compare more than two groups if they showed normal distribution. In case of non-normal distribution, Kruskal-Wallis test followed by Dunn’s multiple comparison post test was conducted. A test was considered significant when p < 0.05. p values are directly shown in graphs. All data are presented as mean ± SEM, and the distribution of the mean value per cell is also shown. For IUE experiments, the number of animals is also specified. Details about sample size and statistical tests are reported in each main and supplementary figure.

## DATA AVAILABILITY

All data needed to evaluate the conclusions in the paper are present in the paper and/or the supplementary materials. Any additional information can be provided by the corresponding author pending scientific review and a completed material transfer agreement. This paper does not report original codes. Further information and requests for resources and reagents should be directed to and will be fulfilled by the corresponding author, Matteo Fossati.

## AUTHOR CONTRIBUTIONS

Conceptualization and writing – original draft: M.B. and M.F.; Formal analysis and data curation: M.B., F.B., E.F., D.P. and M.F; Investigation: M.B., F.B., C.O., M.M., I.R., E.F., M.E., A.F. and M.F.; Methodology and visualization: M.B., F.B. and M.F.; Writing – review & editing: M.B., A.F., D.P. and M.F; Project administration, supervision and funding acquisition: M.F.

## ACKNOWLEDGMENTS

We thank all the members of Fossati’s lab (Humanitas Research Hospital and IN-CNR), Michela Matteoli, Nica Borgese and Sara Francesca Colombo for critical reading of the manuscript and helpful discussions. We thank the Imaging Unit and Animal Facility of Humanitas Research Hospital for their support. This work was supported by the Ministry of Health (Ricerca Finalizzata GR-2018-12366478 to M.F.); Telethon Foundation (GGP20127 and GJC22059 to M.F.); Ministry of Research (PRIN 2022 PNRR P20228FL2Z to M.F.), Angelman Syndrome Alliance (ASA to M.F.) and the Italian Organization for Angelman Syndrome (OR.S.A. to MB).

