## Supplementary figures for "SUBCELLULAR FUNCTIONS OF *UBE3A* ISOFORMS DRIVE SYNAPTIC DYSFUNCTION IN ANGELMAN SYNDROME"

### SUPPLEMENTARY INFORMATION

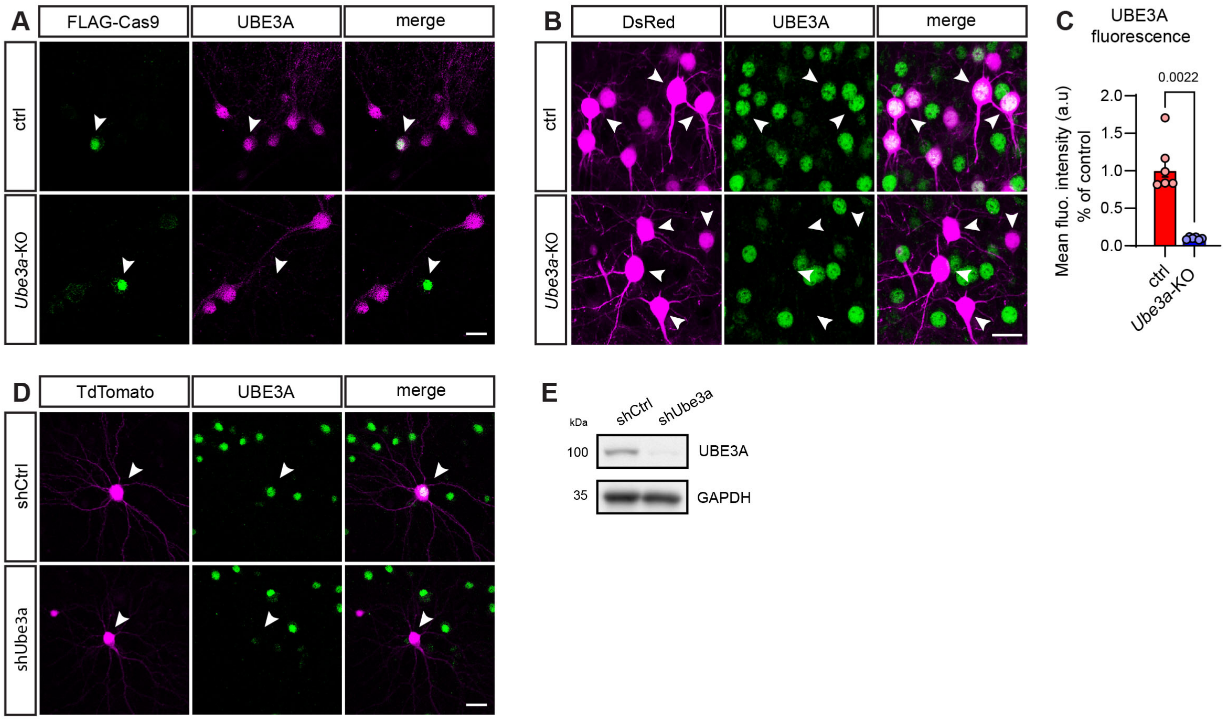


**Figure S1. Validation of CRISPR/Cas9- and shRNA-based approaches to inactivate *Ube3a* expression (related to Fig 1).**

**(A)** Representative confocal images illustrating the effect of CRISPR/Cas9-mediated *Ube3a* knock-out (KO) on UBE3A protein expression in primary cortical neurons transfected at DIV7 and processed for immunocytochemistry at DIV18. Arrowheads point to transfected cells. Scale bar: 20µm. **(B-C)** Representative confocal images (B) and quantification (C) illustrating the effect of CRISPR/Cas9-mediated *Ube3a*-KO on UBE3A protein expression upon IUE at E15.5. Mice were sacrificed and processed for immunohistochemistry at P21. Arrowheads in B point to electroporated neurons and show the efficient suppression of UBE3A expression in *Ube3a*-KO brains. Scale bar: 20 µm. $n_{ctrl}$ = 6 (2); $n_{Ube3a\text{-}KO}$ = 6 (2) brain slices. **(D)** Representative confocal images illustrating UBE3A protein expression in primary culture of cortical neurons at DIV18 and transfected with shRNA Control (shCtrl) or against mouse *Ube3a* (shUbe3a). Arrowheads point to transfected neurons. Scale bar: 20 µm. **(E)** Representative western blot of UBE3A expression in DIV17 cortical neurons transduced with lentiviral vectors driving the expression of shCtrl or shUbe3a. GAPDH is used as loading control. Statistics: Numbers in parentheses indicate the number of animals. Bars indicate mean ± SEM. p values are indicated in the graphs. Mann-Whitney test.

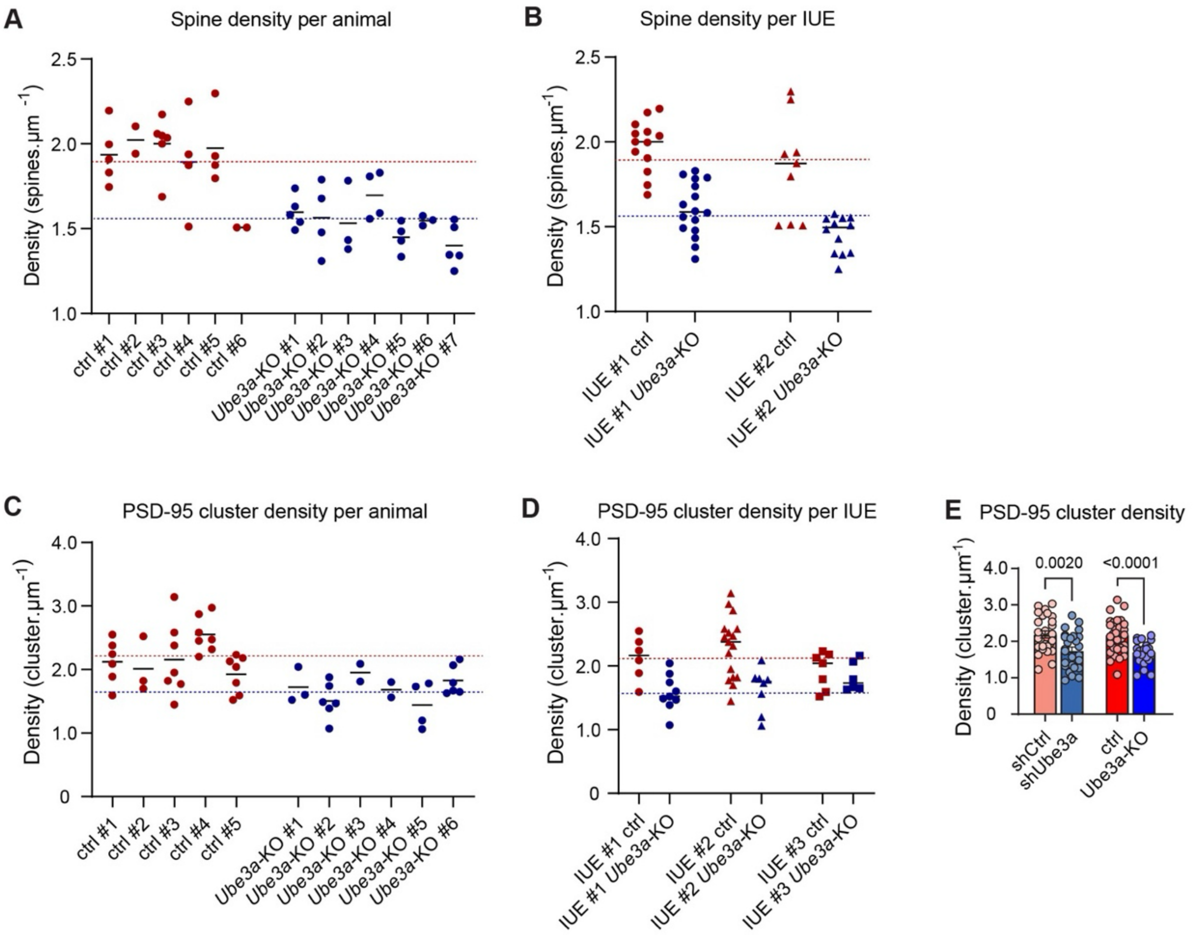


**Figure S2. Effects of *Ube3a*-KO on dendritic spine and PSD-95 cluster density per animal and per IUE (related to Fig 1).**

**(A, C)** Plot showing dendritic spine (A) and PSD-95 cluster (C) density per animal in in layer 2/3 pyramidal neurons expressing CRISPR/Cas9 control (ctrl) and against *Ube3a* (*Ube3a*-KO) at P21 (same data as in Figure 1). Each dot represents one dendrite, and the bar indicates the mean value. Colored dotted lines are the average density of spine density of the corresponding condition (same data of Figure 1). #: mouse identification. **(B, D)** Plot showing dendritic spine (A) and PSD-95 cluster (C) per IUE in in layer 2/3 pyramidal neurons expressing CRISPR/Cas9 control (ctrl) and against *Ube3a* (*Ube3a*-KO) at P21 (same data as in Figure 1). Each dot represents one dendrite, and the bar indicates the mean value. Colored dotted lines are the average density of spine density of the corresponding condition (same data of Figure 1). **(E)** Histograms showing the comparison of PSD-95 cluster density between in utero electroporated layer 2/3 neurons expressing either CRISPR/Cas9 or shRNAs to inactivate Ube3a expression. $n_{ctrl}$ = 31 (5); $n_{Ube3a\text{-}KO}$ = 24 (6); $n_{shCtrl}$ = 27 (7); $n_{shUbe3a}$ = 26 (5). Statistics: bars indicate mean ± SEM. Numbers in parentheses indicate the number of animals. p values are indicated in the graphs. Student t-test.

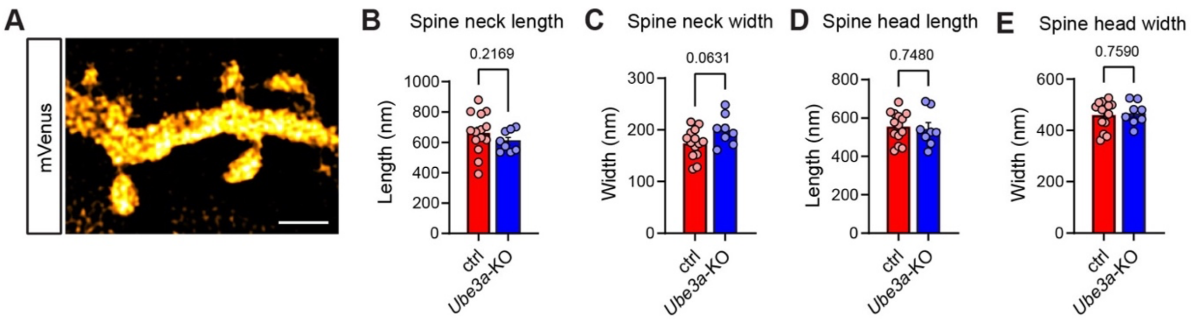


**Figure S3. Analysis of spine morphology using STED microscopy (related to Fig 1).**
**(A)** Representative STED image of a dendrite segment showing spine morphology at high resolution. Scale bar: 1 µm. **(B-E)** Quantification of neck length (B) and width (C), head length (D) and width (E) of dendritic spines in layer 2/3 cortical neurons at P21 expressing CRISPR/Cas9 control (ctrl) or against *Ube3a* (*Ube3a*-KO) using STED microscopy. $n_{ctrl}$ = 15 (6); $n_{Ube3a\text{-}KO}$ = 8 (3). Statistics: bars indicate mean ± SEM. Numbers in parentheses indicate the number of animals. p values are indicated in the graphs. Student t-test and Mann-Whitney test.

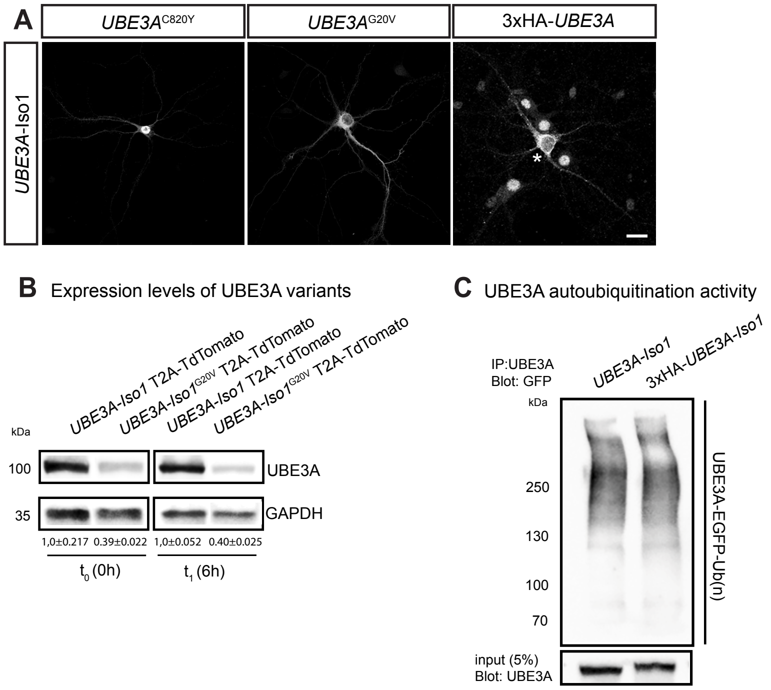


**Figure S4. Subcellular localization, expression levels and enzymatic activity of UBE3A mutants (related to Fig 6).**

**(A)** Representative confocal images illustrating the effect of C820Y, G20V mutations or 3xHA insertion on UBE3A-Iso1 localization. Primary cultures of cortical neurons were transfected at DIV7 with the indicated *hUBE3A-Iso1* variants, fixed and stained at DIV18. Asterisk displays a transfected neuron. Since 3xHA-UBE3A was visualized using an anti-UBE3A antibody, background nuclear signal derives from endogenous UBE3A staining of non-transfected neurons. Scale bar: 20 µm. **(B)** Protein expression of the G20V UBE3A-Iso1 variant. HEK293T cells transfected with indicated plasmids expressing WT or G20V UBE3A-Iso1. Forty-eight hours post-transfection, cells were either harvested ($t_0$) or treated with cycloheximide (CHX, 50 µg/ml) for 6h ($t_6$) followed by western blot analysis. GAPDH was used as loading control. **(C)** Representative western blot showing comparable catalytic activity between WT or 3xHA-tagged UBE3A-Iso1. HEK293T cells were co-transfected with indicated plasmids driving the expression of WT or 3xHA-UBE3A-Iso1 together with EGFP-tagged ubiquitin. Forty-eight hours post-transfection, cells were treated with the proteasome inhibitor MG-132 (30 µM) for 2h and harvested. Protein lysates were immunoprecipitated with an anti-UBE3A antibody then blotted using an anti-GFP antibody to measure UBE3A autoubiquitination (top blot). An anti-UBE3A antibody was used to visualize expression of WT or 3xHA-UBE3A Iso1 (input, bottom blot).
